# Drosophila beta-Tubulin 97EF is upregulated at low temperature and stabilizes microtubules

**DOI:** 10.1101/185827

**Authors:** Faina Myachina, Fritz Bosshardt, Johannes Bischof, Moritz Kirschmann, Christian F. Lehner

## Abstract

**Summary statement:** Ectotherms thrive within an often remarkable temperature range. At low temperature, *betaTub97EF*, a beta-tubulin paralog stabilizing microtubules, is upregulated in a tissue-specific manner in the fly *Drosophila melanogaster*.

**Abstract:** Cells in ectotherms function normally within an often wide temperature range. As temperature dependence is not uniform across all the distinct biological processes, acclimation presumably requires complex regulation. The molecular mechanisms coping with the disruptive effects of temperature variation are still poorly understood. Interestingly, one of five different beta-tubulin paralogs, *betaTub97EF*, was among the genes up-regulated at low temperature in cultured *Drosophila* cells. As microtubules are known to be cold-sensitive, we analyzed whether *betaTub97EF* protects microtubules at low temperatures. During development at the optimal temperature (25°C), *betaTub97EF* was expressed in a tissue-specific pattern primarily in the gut. There, as well as in hemocytes, expression was increased at low temperature (14°C). While *betaTub97EF* mutants were viable and fertile at 25°C, their sensitivity within the well-tolerated range was slightly enhanced during embryogenesis specifically at low temperatures. Changing beta-tubulin isoform ratios in hemocytes demonstrated that beta-Tubulin 97EF has a pronounced microtubule stabilizing effect. Moreover, *betaTub97EF* is required for normal microtubule stability in the gut. These results suggest that *betaTub97EF* up-regulation at low temperature contributes to acclimation by stabilizing microtubules.

## Introduction

Many ecological niches are exposed to ambient temperature fluctuations. Distinct strategies have evolved to cope with the disruptive effects of temperature changes on cellular homeostasis. Endotherms like humans rely primarily on internally generated heat for maintenance of a relatively high and constant core body temperature. However, the majority of organisms are ectotherms. Their internal temperature is primarily dictated by the environment. But even in endotherms, the exposed peripheral cells need to function over a range of temperatures.

Arguably the most extensive analyses of the cellular response to low temperature in eukaryotes so far have been done with yeast (Gasch et al., 2000; Taymaz-Nikerel et al., 2016), confirming the general notion that rapid temperature shifts to nonlethal low temperature disrupt cellular homeostasis, triggering an initial transient stress response followed by persistent acclimation. Moreover, the response to low temperature is quite distinct from the general stress response induced by many other perturbations.

In animals, low temperature acclimation includes additional response levels (Denlinger and Lee, 2010; Hayward et al., 2007). Organ systems govern complex behavioral and metabolic responses. At the cellular level, microtubules have long been recognized to be cold-sensitive in mammals (Correia and Williams, 1983). Microtubules are dynamic cytoskeletal elements involved in innumerable processes (Nogales and Zhang, 2016). They are polymerized from heterodimers of alpha- and beta-tubulin. Heterodimers can associate longitudinally into protofilaments. Lateral interactions between parallel protofilaments organize these into hollow cylindrical tubes. Both alpha- and beta-tubulin bind GTP at a conserved site. GTP-tubulin heterodimers incorporate efficiently at growing microtubule ends. Polymerization is coupled to hydrolysis of the GTP bound to beta-tubulin. This regulates microtubule dynamics and gives rise to dynamic instability where growing phases alternate with shrinking phases (Mitchison and Kirschner, 1984; Zhang et al., 2015). Above a critical concentration, GTP-tubulin purified from mammalian sources polymerizes into microtubules at 37°C, largely driven by the entropically favorable displacement of structured water at the subunit interfaces (Correia and Williams, 1983). Reflecting the positive correlation between entropic drive and temperature, these microtubules depolymerize after a shift to 4°C in a GTP hydrolysis-independent manner (Fygenson et al., 1994).

Psychrophilic ectotherms assemble microtubules even below 0°C. Analysis of their genes have revealed amino acid changes that likely alter the temperature dependence of microtubule stability (Chiappori et al., 2012; Detrich et al., 2000; Modig et al., 2000; Tartaglia and Shain, 2008). While such evolutionary adaptations support life at constant low temperature, organisms exposed to short term thermal fluctuations presumably depend on temperature controlled regulation of microtubule dynamics. In principle, regulated expression of tubulin paralogs with distinct properties is conceivable, as well as control by posttranslational modifications and indirectly via microtubule-associated proteins (MAPs) and motors. Paralogs encoding distinct alpha- and beta-tubulins have evolved in many lineages (Findeisen et al., 2014). Sequence differences are found predominantly in the C-terminal tails, where also most posttranslational modifications occur. These modifications are thought to generate a ‘tubulin-code’ controlling binding and function of many MAPs and motors (Gadadhar et al., 2017; Janke, 2014; Sirajuddin et al., 2014; Yu et al., 2015).

In mammals, MAP6 adapts its conformation according to temperature to maintain microtubule networks in cells exposed low temperature (Delphin et al., 2012). Moreover, the recent production of pure tubulin isoforms has definitely established inherent isoform-specific differences in microtubule dynamics *in vitro* (Pamula et al., 2016; Ti et al., 2016; Vemu et al., 2016).

Genetic analyses in *Drosophila melanogaster* have provided the first unequivocal evidence for paralog-specific functions *in vivo* (Ludueña and Banerjee, 2008). The genome of *D. melanogaster* contains five alpha-tubulin genes, as well as five beta-tubulin genes (*betaTub56D*, *betaTub60D*, *betaTub85D*, *betaTub97EF*, and *CG32396* designated here as *betaTub65B*). The most divergent beta-tubulin paralogs (*betaTub85D* and *betaTub65B*) are expressed exclusively in testis. While *betaTub65B* has not yet been studied, *betaTub85D* has been characterized in considerable detail. It is required in the germline for male meiotic divisions and sperm axoneme formation (Fuller et al., 1988; Kemphues et al., 1982). These processes still fail when *betaTub60D*, which is not normally expressed in the male germline, is expressed instead of *betaTub85D* (Hoyle and Raff, 1990). The normal expression of *betaTub60D* is also tissue-specific, but considerably more complex than that of the testis-specific *betaTub85D*. During embryogenesis *betaTub60D* expression starts in differentiating mesodermal cell types, but occurs also in chordotonal organs, imaginal discs and somatic cells of the adult gonads (Kimble et al., 1990; Rudolf et al., 2012). The zygotic expression of *betaTub56D* has also pronounced tissue-specificity, but at the start of embryogenesis the abundant maternal *betaTub56D* contribution is the only source of beta-tubulin (Buttgereit et al., 1996).

Here we report that *betaTub97EF*, an uncharacterized beta tubulin paralog, is upregulated at low temperature in specific cell types. Moreover, we demonstrate that it increases microtubule stability.

## Results

### *betaTub97EF* encodes a temperature-regulated beta-tubulin

Genome-wide expression profiling with *Drosophila* S2R+ cells over a range of different growth temperatures revealed *betaTub97EF* among the temperature-responsive genes. Compared to 25°C, the presumed optimal temperature, *betaTub97EF* transcript levels were increased at lower temperatures (11 and 14°C) and decreased at a higher temperature (30°C) in multiple independent experiments (Fig. 1A; data not shown). In another cell line, embryonic cells immortalized exploiting *Act5c>Ras85D*^*G12V*^ as described recently (Simcox et al., 2008), *betaTub97EF* transcript levels were also twofold higher at 14°C compared to 30°C. In contrast to *betaTub97EF*, other tubulin genes were at most marginally affected by growth temperature (Fig. 1A). Independent analyses with quantitative reverse transcription polymerase chain reaction (qRT-PCR) confirmed the microarray results (Fig. 1B,C). According to qRT-PCR, *betaTub97EF* transcripts amount to about 8% of the total beta-tubulin transcripts, while *betaTub56D* and *betaTub60D* contribute 46% each in S2R+ cells at 25°C (Fig. 1B). At 14°C, *betaTub97EF* transcripts are more than twofold higher (Fig. 1C).

**Fig. 1.**
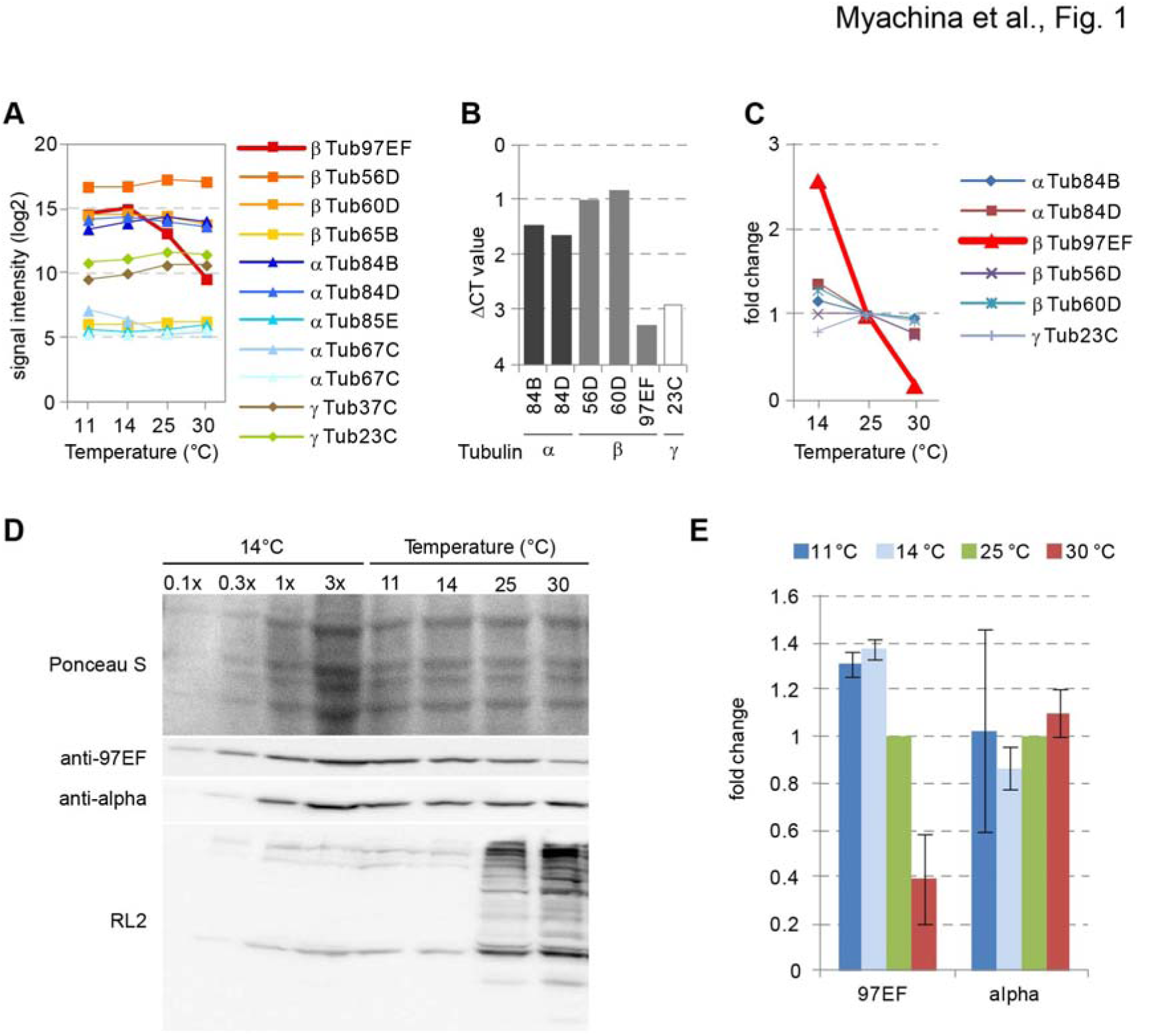
*betaTub97EF* expression is induced at low temperature. The effect of growth temperature on transcript and protein levels of *Drosophila* tubulin genes in S2R+ cells was analyzed with microarrays (A), qRT-PCR (B,C) or immunoblotting (D,E). (A) Temperature dependence of signal intensities on microarray probes. (B) Transcript levels of significantly expressed tubulin genes at 25°C. ΔCT values are indicated. A ΔCT value of zero indicates a transcript level equal to average expression of the genes used for normalization; higher ΔCT values indicate a relative decrease in expression. (C) Temperature dependence of transcript levels of significantly expressed tubulin genes. For each gene, the level at 25°C was set to 1. (D) Total extracts of S2R+ cells grown at 11, 14, 25 or 30°C were analyzed after western blotting by Ponceau S and probing with the indicated antibodies. RL2 detects O-GlcNAc modification on proteins which is tightly correlated with growth temperature (Radermacher et al., 2014). Equal amounts of protein were loaded except for the first four lanes covering a dilution series of the 14°C extract as internal reference for quantification. (E) Signals obtained with anti-betaTubulin97EF and anti-alpha-tubulin from immunoblots as shown in (D) were quantified. Ponceau S staining intensity was used for normalization. Signal intensities observed at 25°C were set to 1. Bars indicate average (+/- s.d.; n = 3).

To study temperature effects on protein abundance, we raised an antibody against beta-Tubulin 97EF using a C-terminal peptide for immunization. The C-terminal tails are the most divergent regions of the highly conserved beta-tubulins (Fig. S1). They have previously permitted production of paralog-specific antibodies (Kimble et al., 1989; Leiss et al., 1988). Immunoblotting experiments with extracts from bacteria expressing GST fusion proteins with extensions identical to the tails of the different beta-tubulin paralogs were used for initial antibody characterization (Fig. S2A,B). Immunoblotting with these extracts also demonstrated that the mouse monoclonal antibody E7 (Chu and Klymkowsky, 1989) detects with almost equal sensitivity all of the beta-tubulin paralogs, except beta-Tubulin 65B, a most divergent uncharacterized paralog with low testis-specific expression. The specificity of the newly generated antibodies for beta-Tubulin 97EF was further confirmed by immunoblotting and immunolabeling experiments using *betaTub97EF* mutants (see below). Quantitative immunoblotting indicated that the amount of beta-Tubulin 97EF in S2R+ cells is only about 2% of the total beta tubulin protein at 25°C (Fig. S1C). However, immunoblotting with extracts of S2R+ cells grown at different temperatures demonstrated that beta-Tubulin 97EF levels were temperature-dependent. Although slightly less pronounced than *betaTub97EF* transcripts, protein was significantly more abundant at low temperature (11 and 14°C) and reduced at high temperature (30°C) compared to 25°C (Fig. 1D,E). As expected (Radermacher et al., 2014), control immunoblotting with RL2, an antibody recognizing O-GlcNAc modified proteins, revealed the opposite temperature dependence. In conclusion, in cultured cells both mRNA and protein levels of *betaTub97EF* respond to temperature. Higher levels are present at low temperatures.

### *betaTub97EF* expression occurs predominantly in the gut

Three of the five beta-tubulin paralogs, *betaTub56D*, *betaTub60D* and *betaTub85D* have been studied in considerable detail (Rudolf et al., 2012, and references given therein) but not *betaTub97EF*. Therefore, the pattern of *betaTub97EF* expression during development at 25°C was analyzed. By immunoblotting (Fig. 2A), beta-Tubulin 97EF could not be detected at the onset of embryonic development, consistent with RNA-Seq data (Graveley et al., 2011). Zygotic expression started between 6-9 hours after egg deposition (AED) and increased during the second half of embryogenesis. The specificity of the corresponding immunoblot signals (Fig. 2A) was confirmed with extracts of *betaTub97EF* mutant embryos (see below).

**Fig. 2.**
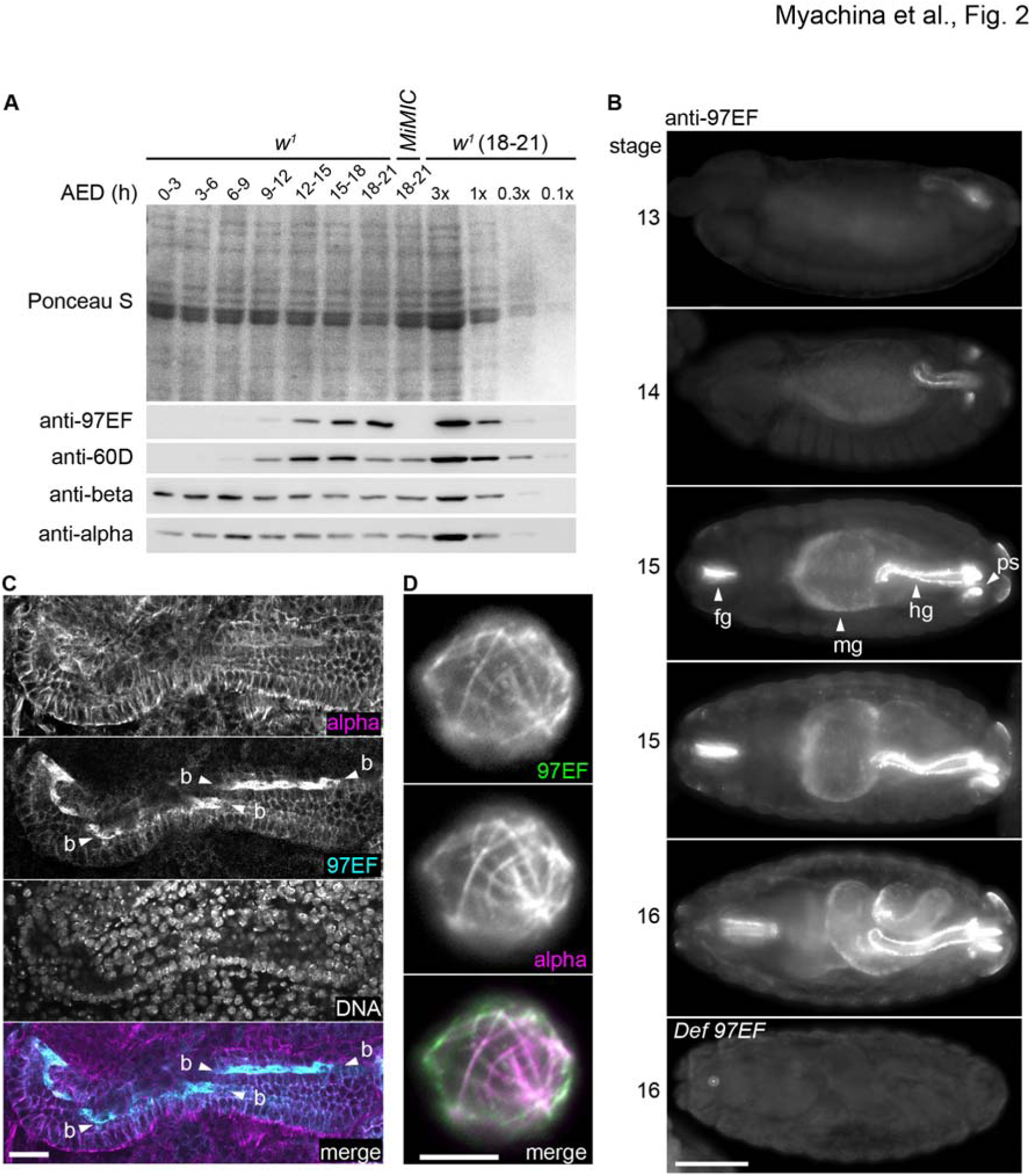
*betaTub97EF* expression during embryogenesis. (A) Total extracts from *w*^*1*^ embryos at different ages AED. Equal numbers of embryos at each age interval were loaded apart from a dilution series. An extract from *betaTub97EF* null mutants (*MiMIC*) was included. Ponceau S staining was used as a control for equal loading. Replicate immunoblots were probed with the indicated antibodies. (B) *w*^*1*^ embryos were stained with anti-beta-Tubulin 97EF and a DNA stain (not shown). In addition, an embryo homozygous for *Df(3R)BSC460* which deletes *betaTub97EF* is shown in the bottom panel (*Def 97EF*). Numbers on the left side indicate the displayed stages. Arrowheads indicate foregut (fg), midgut (mg), hindgut (hg) and posterior spiracles (ps). (C) Hindgut region (13-15 hours AED) labeled as indicated. Boundary cells (bc) are pointed out by arrowheads. (D) Larval hemocyte after double labeling with anti-beta-Tubulin 97EF and anti-alpha-tubulin. Scale bar 50 μm (B), 10 μm (C) and 5 μm (D).

For comparison, immunoblotting with anti-beta-Tubulin 60D confirmed an expression program similar to beta-Tubulin 97EF with a slightly earlier onset and peak of zygotic expression (Fig. 2A). Moreover, anti-pan-beta-tubulin E7 revealed maximal levels of total beta-tubulin at the start of embryogenesis (Fig. 2A) when the abundant maternally contributed beta-Tubulin 56D is present (Rudolf et al., 2012). The monoclonal antibody DM1A (Blose et al., 1984) was used to detect alpha-tubulins (Fig. 2A). Based on its epitope characterization (Breitling and Little, 1986), DM1A presumably reacts with all or most of the *Drosophila* alpha-tubulins but the precise affinities for the different isoforms are not known.

Immunofluorescent labeling was performed to determine the spatial pattern of beta-Tubulin 97EF expression during embryogenesis (Fig. 2B). beta-Tubulin 97EF was not detected until stage 12 when weak specific signals were observed in the developing hindgut. Subsequently signal intensities increased in the hindgut. Expression was also detected in foregut and posterior spiracles, as well as more weakly in the midgut (Fig. 2B). The predominantly gut-specific expression pattern of beta-Tubulin 97EF is clearly distinct from that of beta-Tubulin 60D with its primarily muscle-specific expression pattern (Kimble et al., 1989; Leiss et al., 1988), and data not shown). The maternally contributed, abundant beta-Tubulin 56D is ubiquitously and uniformly distributed during embryogenesis, as indicated by E7 staining (data not shown).

To characterize anti-beta-Tubulin 97EF staining in further detail, the region with the embryonic hindgut was analyzed at high magnification (Fig. 2C). This revealed beta-Tubulin 97EF in the epithelial cells of the hindgut, but not in the surrounding visceral muscles. Anti-beta-Tubulin 60D resulted in the opposite pattern (data not shown). Within the hindgut epithelium, beta-Tubulin 97EF was most abundant in the so-called boundary cells (Fig. 2C), a row of single cell width extending along both sides of the hindgut tube between the ventral and dorsal compartment (Iwaki and Lengyel, 2002).

At maximal resolution, near perfect spatial overlap of the signals obtained by double labeling with anti-beta-Tubulin 97EF and anti-alpha-tubulin was observed (Fig. 2D). For these high resolution analyses, we used released larval hemocytes which spread out flat on coverslips, providing superior optical conditions. Although beta-Tubulin 97EF was colocalized with alpha-tubulin, the spatial intensity distributions of the signals obtained by double labeling with the two antibodies were not identical. Accordingly, beta-Tubulin 97EF might be enriched in cortical microtubules, but alternative explanations are not excluded (see discussion).

Immunolabeling revealed that beta-Tubulin 97EF is also present in the larval and adult gut (Fig. S3B,C). Signals in enterocytes were rather uniform, i.e., not as in the embryonic hindgut where boundary cells have higher levels. beta-Tubulin 97EF was not only detected in the gut. Specific signals, although weaker, were observed in wing imaginal discs, larval hemocytes and ovarial follicle cells (see below and data not shown).

We conclude that *betaTub97EF* expression occurs in a pattern distinct from that of other beta-tubulin paralogs. *In situ* hybridization and a *betaTub97EF>GFP* reporter allele suggested that tissue and cell-type specificity of *betaTub97EF* expression is transcriptionally controlled (data not shown).

### *betaTub97EF* upregulation by low temperature is tissue-specific

To address the physiological role of *betaTub97EF*, we characterized mutant alleles (Fig. 3A). Three *Minos* transposon insertions in the *betaTub97EF* region were analyzed (Fig. 3A). PCR assays confirmed the annotated insertion positions. The insertion *Mi{MIC}betaTub97EF^MI06334^* within the first intron (*betaTub97EF*^*MiMIC*^ in the following) introduces a splice acceptor site followed by translational stops in all three reading frames. By conversion of this insertion (Venken et al., 2011), a derivative with an extra EGFP-encoding exon within this first intron was obtained (*betaTub97EF*^*EGFP*^). The predicted protein product is expected to be non-functional because the EGFP insertion disrupts the tubulin core domain. Indeed, beta-Tubulin97EF^EGFP^ was not incorporated into microtubules according to fluorescence microscopy.

**Fig. 3.**
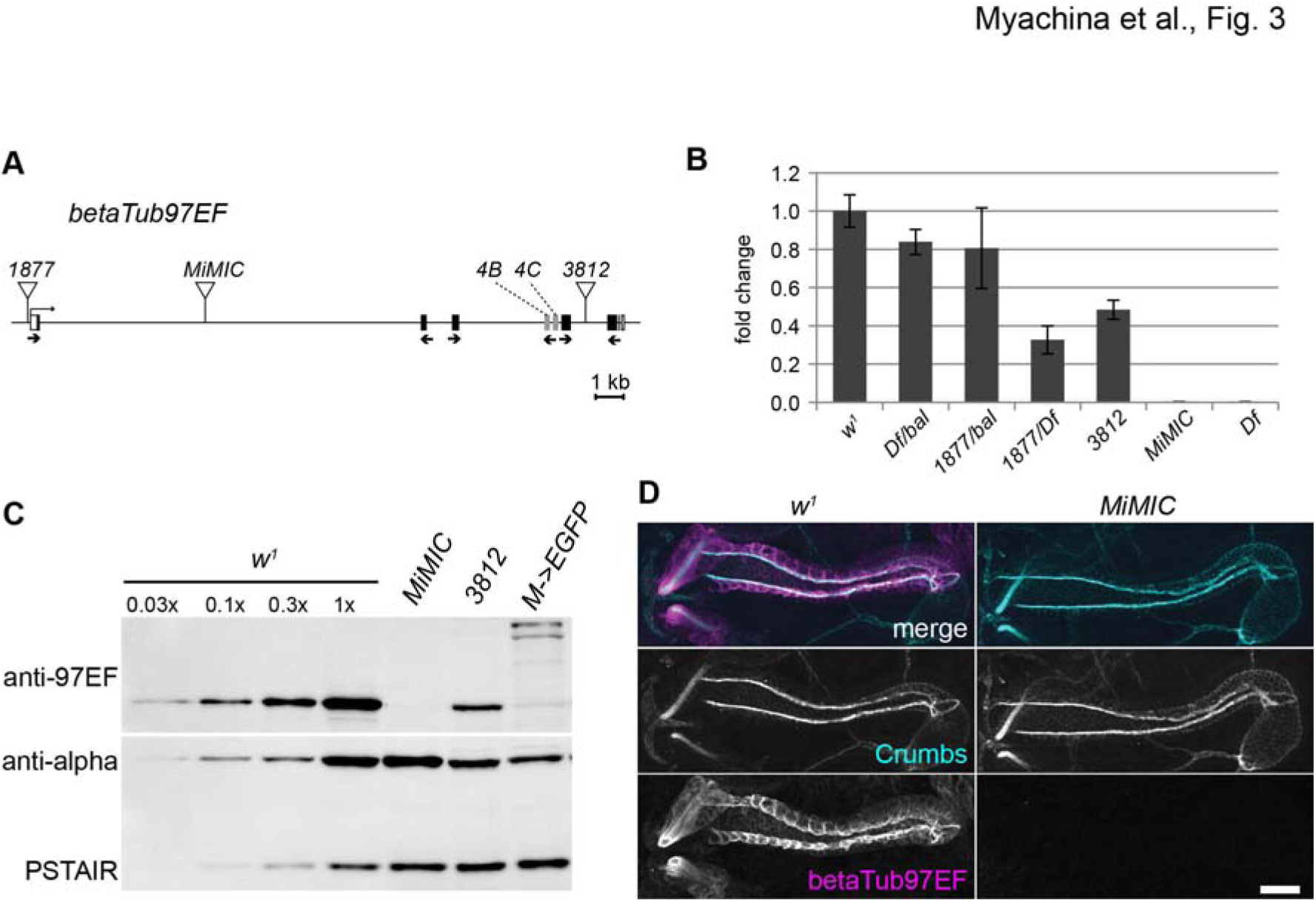
Characterization of mutant *betaTub97EF* alleles. (A) Scheme of the *betaTub97EF* region. Transposon insertions (*1877*, *MiMIC* and *3812*) are indicated by triangles and exons by boxes with black filling in coding region, except for the two mutually exclusive exons 4B and 4C which are shown in grey. Small arrows below exons indicate the three primer pairs used for qRT-PCR. (B) *betaTub97EF* transcript levels were determined by qRT-PCR with embryos of the indicated genotypes (*Df* = *Df(3R)BSC460*). Results with the three primer pairs were highly concordant in a given genotype and were therefore averaged. Transcript levels are indicated by bars (average +/- s.d., n = 3 primer pairs). Those of *w*^*1*^ control embryos were set to 1. (C) Total extracts from embryos (15-16 hours AED) analyzed by immunoblotting with genotypes and antibodies indicated. Anti-alpha-tubulin and anti-PSTAIR were used as a mixture to probe the replicate immunoblot shown in the lower panel. (D) Hindgut region in embryos (14-15 hours AED) after immunostaining with genotypes and antibodies as indicated. Crumbs is maximally expressed in the boundary cells of the hindgut. Scale bar 20 μm.

By qRT-PCR (Fig. 3B) and immunoblotting (Fig. 3C), we analyzed how *betaTub97EF* expression was affected by the transposon insertions. While two, *Mi{ET1}betaTub97EF*^*MB01877*^ and *Mi{ET1}betaTub97EF*^*MB03812*^, reduced expression, *betaTub97EF*^*MiMIC*^ appeared to abolish it completely (Fig. 3B,C). Residual beta-Tubulin 97EF, potentially expressed from the *betaTub97EF*^*MiMIC*^ allele, cannot be more than 3% of the normal level, because stronger expression would have been detectable (Fig. 3C). In case of *betaTub97EF*^*EGFP*^, immunoblotting confirmed the expected expression of a larger beta-Tubulin 97EF variant with an internal EGFP domain (Fig. 3C). As anti-beta-Tubulin 97EF also revealed a weak band at the position of normal beta-Tubulin 97EF, rare skipping of the extra EGFP exon present in the *betaTub97EF*^*EGFP*^ allele cannot be excluded and might allow expression of normal beta-Tubulin 97EF up to 10% of wild-type levels. Alternatively, the weak band might represent one of the obvious beta-Tubulin 97EF^EGFP^ degradation products, rather than residual normal beta-Tubulin 97EF.

Unexpectedly given the conservation of the *betaTub97EF* ortholog during an estimated 300 million years of insect evolution (Findeisen et al., 2014), *betaTub97EF* function was found to be dispensable for viability and fertility under standard laboratory conditions. *betaTub97EF*^*MiMIC*^ could be propagated as a homozygous stock. Compared to *w*^*1*^, fertility was 25% lower in both sexes in flies with *betaTub97EF*^*MiMIC*^ over a deficiency deleting *betaTub97EF* (*Df(3R)BSC460*).

Since *betaTub97EF* expression is especially high in boundary cells of the embryonic hindgut, we analyzed whether these cells are present in *betaTub97EF*^*MiMIC*^ mutants. Labeling of Crumbs, an apical membrane protein expressed at very high levels in boundary cells (Kumichel and Knust, 2014), demonstrated that these cells are present in *betaTub97EF*^*MiMIC*^ mutants (Fig. 3D). To compare hindgut morphology in *betaTub97EF*^*MiMIC*^ mutants and control embryos with ultrastructural resolution, we applied focused ion beam scanning electron microscopy (FIBSEM). While this failed to expose abnormalities in *betaTub97EF*^*MiMIC*^ mutants, our three-dimensional reconstructions revealed that boundary cells differentiate striking undulae propagating longitudinally from cell to cell along the hindgut epithelial tube (Fig. S4), rather than prominent microvilli as suggested previously based on two-dimensional EM analyses (Iwaki and Lengyel, 2002; Kumichel and Knust, 2014; Soplop et al., 2012).

The hindgut is one of the organs that develops left/right asymmetry during development (Coutelis et al., 2014). *betaTub97EF*^*MiMIC*^ mutants (n = 100) were found to have normal left/right asymmetry in larval and adult hindgut. We also addressed whether absence of beta-Tubulin 97EF from the gut might increase sensitivity to hyperosmotic diet. Therefore, survival to the adult stage after larval development in high salt food was analyzed. However, *betaTub97EF*^*MiMIC*^ mutants were not more salt-sensitive than *w*^*1*^ control (data not shown).

Our phenotypic analyses suggested that *betaTub97EF* does not provide crucial functions at temperatures close to the optimum of 25°C. However, since *betaTub97EF* expression in S2R+ cells is increased at low temperature (Fig. 1), this gene might be of a greater importance during development and life at sub-optimal temperatures. First we analyzed whether regulation of *betaTub97EF* expression by temperature occurs also in embryos. After egg collection at 25°C, aliquots were shifted at 3-6 hours AED to either 14, 25 or 30°C. Once embryos had reached stage 16 (peak of *betaTub97EF* expression), total RNA was analyzed by qRT-PCR. Among tubulin paralogs, *betaTub97EF* is unique with regard to the presence of two mutually exclusive exons (4B and 4C) within the region coding for the conserved tubulin protein core (Fig. 3A) (Findeisen et al., 2014). These two mutually exclusive exons appear to be characteristic for dipterans (Findeisen et al., 2014). In *D. melanogaster* they share 77% identity at the amino acid sequence level (Fig. S1). Based on splice junction counts (Graveley et al., 2011), the 4B variant is about 15 fold more abundant than the 4C variant throughout development. Our qRT-PCR analyses were done with two distinct primer pairs designed to detect specifically one or the other isoform. While the minor 4C transcripts were found to be downregulated, the major 4B transcripts were up-regulated at lower temperature (11°C) (Fig. 4A).

**Fig. 4.**
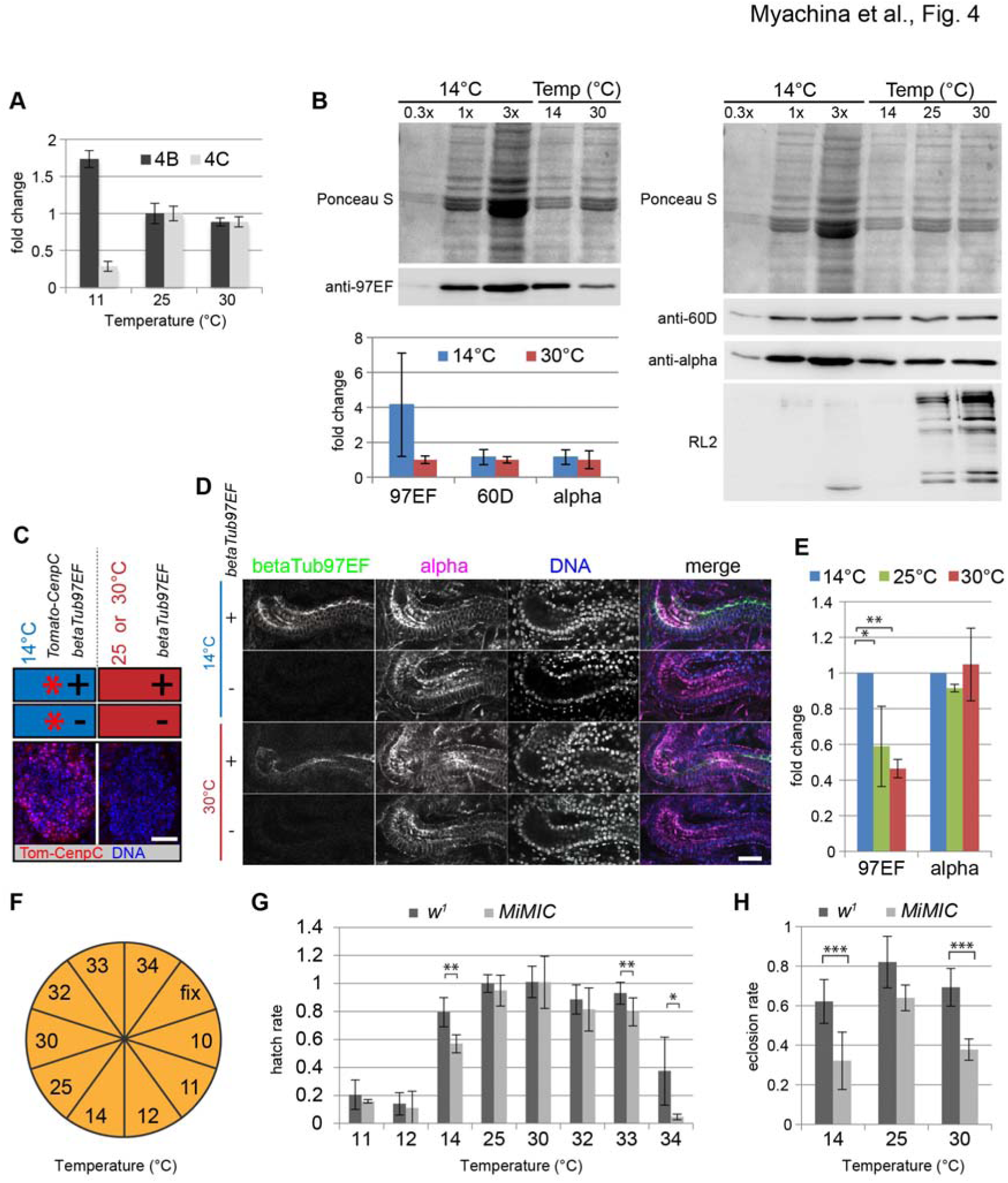
*betaTub97EF* upregulation at low temperature in embryos is tissue specific. (A) qRT-PCR revealed that the major *betaTub97EF* transcript isoform with exon 4B is upregulated at low temperature, while the minor alternative transcript with exon 4C is downregulated. Expression levels at 25°C were set to 1. Bars indicate average (+/- s.d.; n = 3 technical replicates). (B) Immunoblotting revealed that beta-Tubulin 97EF protein is upregulated in embryos at low temperature, in contrast to beta-Tubulin 60D and alpha-tubulin. Bar diagram represents average immunoblot signal intensities at 14° and 30°C (+/- s.d.; n = 3 biological replicates). Ponceau S staining intensity was used for normalization. Intensities observed at 30°C were set to 1. (C-E) Upregulation of beta-Tubulin 97EF at low temperature is tissue-specific. (C) Experimental strategy for accurate quantification of anti-beta-Tubulin 97EF immunofluorescence signals. See text for further explanations. Scale bar 20 μm. (D) Confocal sections through hindgut illustrate labeling with the indicated antibodies and DNA staining in embryos of the indicated genotypes after development at 14°C and 30°C, respectively. Scale bar 20 μm. (E) Quantification of signal intensities obtained with anti-beta-Tubulin 97EF and anti-alpha-tubulin in the hindgut epithelium. Expression levels observed at 14°C were set to 1. Bars indicate average (+/- s.d., n = 6 embryos). (F-H) Temperature sensitivity of *betaTub97EF* mutant development. (F) Experimental strategy for the analyses at different temperatures. Egg collections were divided into aliquots and incubated at the indicated temperatures. One part (fix) was used for the determination of the number of unfertilized and overaged embryos. (G) Rate of larval hatching at different temperatures. (H) Rate of development to the adult stage at different temperatures. Bars indicate average (+/- s.d., n = 3). * p < 0.05, ** p< 0.01, *** p < 0.001 (*t* test).

Aliquots of the same embryos were also analyzed by immunoblotting (Fig. 4B). This confirmed that beta-Tubulin 97EF protein was up-regulated after development at low temperature. On average, beta-Tubulin 97EF levels were fourfold higher at 14°C compared to 30°C (Fig. 4B). Although with variation in extent, higher beta-Tubulin 97EF levels at the lower temperature were consistently observed in each of three experiments. In contrast, signals obtained by immunoblotting with anti-beta-Tubulin 60D and anti-alpha-tubulin were not affected by temperature (Fig. 4B), indicating that induction by low temperature is specific for the *betaTub97EF* paralog.

To address whether the upregulation of beta-Tubulin 97EF at low temperature occurs globally or in a tissue-specific manner we performed quantitative immunofluorescent labeling. To achieve maximal accuracy and minimize problems caused by potential fixation and staining artefacts, these experiments were performed by combining four different types of stage 16 embryos before fixation and staining in a pool (Fig. 4C). Two of these embryo types were processed after development at 14°C (Fig. 4C, blue). To identify these “cold” embryos during the microscopic analyses, they were collected from strains expressing red fluorescent centromere protein Cenp-C (*g-tdTomato-CenpC*) (Fig. 4C, red stars). In contrast, the two other types, the “warm” embryos, processed after development at a higher temperature (25°C or 30°C), did not have this marker transgene. Based on red fluorescent centromere signals (as illustrated with the brain lobe regions in Fig. 4C) “cold” and “warm” embryos were identified. Moreover, in order to quantify the level of non-specific signals resulting from anti-beta-Tubulin 97EF labeling, both the “cold” and the “warm” embryos were a mixture. Half were *betaTub97EF*^*MiMIC*^ mutants and the other half had two wild-type alleles of *betaTub97EF* (Fig. 4C, + and -, respectively). Green fluorescent anti-beta-Tubulin 97EF labeling revealed the *betaTub97EF* genotype during the microscopic analyses. In addition, anti-alpha-tubulin labeling was visualized using a far-red fluorophore.

Quantification of the specific, background-corrected anti-beta-Tubulin 97EF signals indicated that beta-Tubulin 97EF levels were 1.7 fold higher in the embryonic hindgut at 14°C compared to 25°C, and 2 fold higher when compared to 30°C (Fig. 4D,E). Tissue-specific beta-Tubulin 97EF upregulation in the hindgut at low temperature was also observed when *g-tdTomato-CenpC* was used for marking the “warm” instead of the “cold” embryos. In contrast, anti-alpha-tubulin signals were not affected by temperature (Fig. 4D,E). Analogous quantification of signals within epidermis and somatic muscles did not reveal any temperature effects, neither on beta-Tubulin 97EF nor on alpha-tubulin signals (data not shown).

Although low temperature induces higher beta-Tubulin 97EF expression in the hindgut, this organ appeared to be morphologically normal in *betaTub97EF*^*MiMIC*^ mutants even after embryonic development at 14°C or 12°C (data not shown). Despite normal morphology, certain cells or tissues might not be fully functional in *betaTub97EF*^*MiMIC*^ mutants. Therefore we analyzed temperature sensitivity of development to the larval and adult stages. Embryos were collected at 25°C and aliquots were shifted after gastrulation, but before the onset of *betaTub97EF* expression, to temperatures within the range of 10 to 34°C (Fig. 4F). The rate of larval hatching from eggs (Fig. 4G) as well as the rate of development to the adult stage was determined (Fig. 4H). At 10°C, no larvae were observed to hatch, neither from control nor from *betaTub97EF* mutant eggs. Similarly, only very few larvae hatched at 11°C and 12°C (Fig. 4G). Interestingly, compared to control, 25% fewer *betaTub97EF*^*MiMIC*^ mutants completed embryogenesis successfully at 14°C (Fig. 4G). This temperature of 14°C is the lower limit for completion of the entire life cycle for standard *D. melanogaster* laboratory strains. Therefore, *betaTub97EF* function appears to be important at the lower end of the temperature range compatible with the complete life cycle. In contrast at 30°C, the upper end of this compatible range, there were no significant survival differences between *betaTub97EF*^*MiMIC*^ mutants and control, as well as at the optimum temperature of 25°C (Fig. 4G). Only at more drastically elevated temperatures, embryogenesis of *betaTub97EF* mutants was less successful than in controls (Fig. 4G). When analyzing development up to the adult stage, *betaTub97EF* mutants were found to be significantly weaker than control at temperatures below as well as above the optimum (Fig. 4H). By immunoblotting with total larval extracts, we were actually unable to detect beta-Tubulin 97EF upregulation at low temperature at this developmental stage (data not shown). In conclusion, within the temperature range where the full *D. melanogaster* life cycle can be completed with a normal genotype, the importance of *betaTub97EF* function for successful embryogenesis increases towards the lower limit, supporting the notion that *betaTub97EF* upregulation at low temperature contributes to acclimation.

### beta-Tubulin 97EF generates stable microtubules

Functional redundancy of co-expressed beta-tubulin paralogs might explain the mild phenotypic consequences of *betaTub97EF* null mutations. In *betaTub97EF* mutant embryos, the maternally provided abundant beta-Tubulin 56D (Buttgereit et al., 1991) is present at least initially. To address potential functional redundancy, we first characterized *betaTub56D* mutant alleles. We studied two transposon insertions, *betaTub56D*^*NP0949*^ and *betaTub56D*^*EY12330*^, that appeared likely to be detrimental for *betaTub56D* function (Fig. 5A). Both insertions were found to be homozygous lethal, as well as over a deficiency (*Df(2R)BSC782*) that eliminates 5’ upstream sequences including the promoter region and also the first *betaTub56D* exon (Fig. 5A). Moreover, the insertions failed to complement each other. Finally, the developmental lethality associated with *betaTub56D*^*NP0949*^/*Df(2R)BSC782* was completely suppressed by *UASt-betaTub56D* expression driven by *alphaTub84BP-GAL4* (Fig. 5A). Therefore, *betaTub56D*^*NP0949*^ and *betaTub56D*^*EY12330*^ are loss of function alleles of *betaTub56D*.

**Fig. 5.**
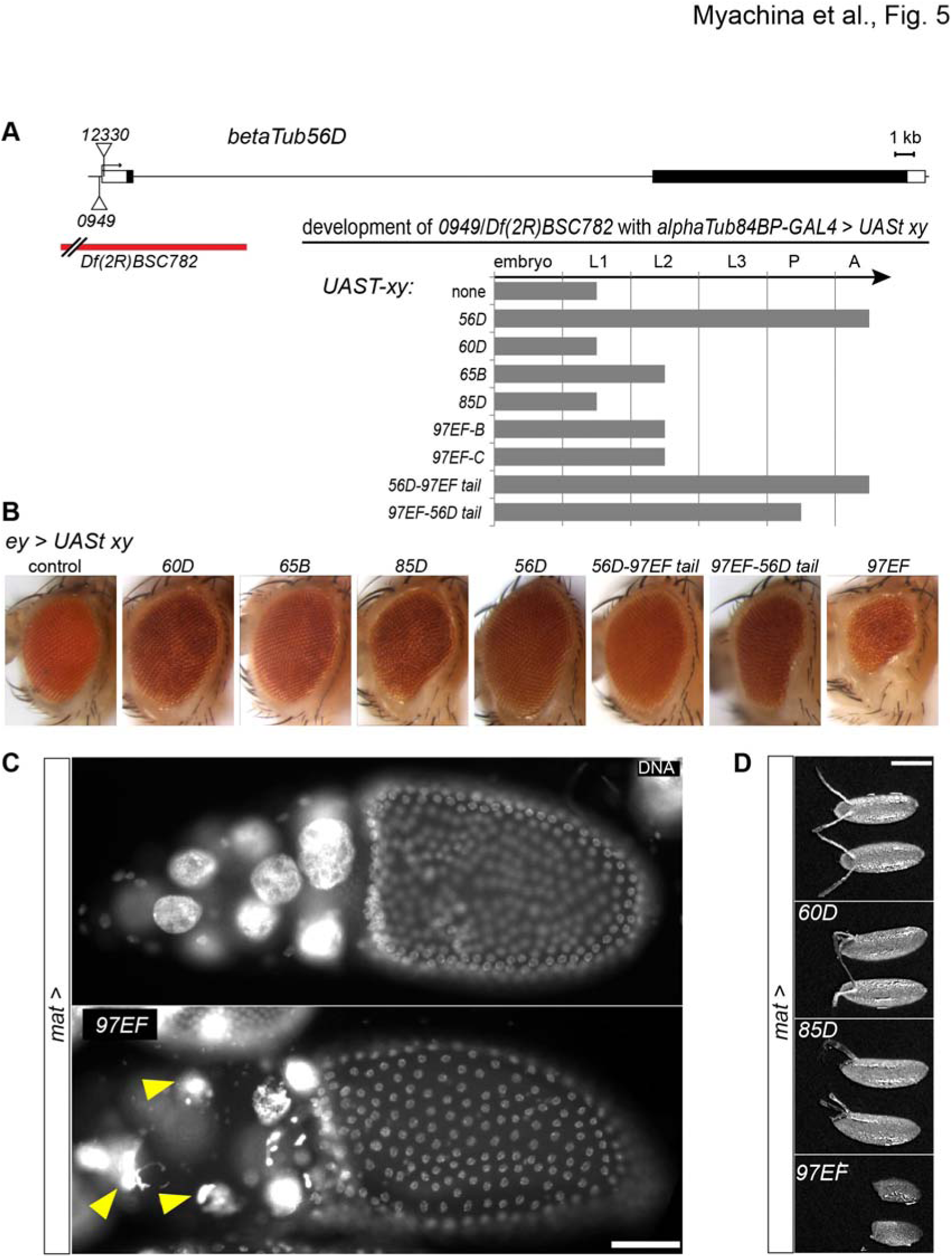
Functional specialization of beta-Tubulin 97EF. Functional specialization was addressed by complementation experiments involving expression of beta-tubulin paralogs in *betaTub56D* mutants (A) as well as after ectopic expression (B,C). (A) The P-element insertions *NP0949* and *EY12330* within the *betaTub56D* region are indicated (triangles), as well as the intragenic breakpoint of the deficiency *Df(2R)BSC782*. Exons are represented by boxes with black filling indicating coding region. Grey bars indicate the extent of development of *betaTub56D*^*NP0949*^/*Df(2R)BSC782* mutants with *alphaTub84BP-GAL4* driven expression of the indicated *UASt-beta-tubulin* transgenes. Larval instar stage 1-3 (L1-L3), pupal stage (P) and adult stage (A). (B) Eye phenotype in adults after *ey-GAL4* driven expression of the indicated *UASt-beta-tubulin* transgenes. (C,D) *mat-4-GAL4VP16* was used for expression of the indicated *UASp-beta-tubulin* transgenes during oogenesis. (C) DNA staining in stage 10B egg chambers. Arrowheads point to some of the abnormal nurse cell nuclei. (D) Morphology of deposited eggs. Scale bar 50 μm.

The lethality associated with *betaTub56D*^*NP0949*^/*Df(2R)BSC782* occurred after embryogenesis during the first larval instar (Fig. 5A), suggesting that the maternal *betaTub56D* contribution is sufficient for successful completion of embryogenesis. Indeed, the abundant maternal *betaTub56D* contribution was found to be stable throughout embryogenesis, as revealed by immunolabeling experiments using *Df(2R)BSC782* (data not shown). This maternal *betaTub56D* contribution might explain why hindgut development appeared to be normal even when zygotic function of both *betaTub97EF* and *betaTub56D* were missing during embryogenesis (data not shown).

Paralog retention after gene duplication during evolution usually depends on functional specialization at the level of expression and/or protein function. To address protein specialization, we compared the *betaTub56D* mutant rescue potential of the different beta-tubulin paralogs (Fig. 5A). For paralog expression in *betaTub56D* mutants (*betaTub56D*^*NP0949*^/*Df(2R)BSC782)*, we always used the same *GAL4* driver (*alphaTub84BP-GAL4*) in combination with *UASt* transgenes that differed only within the coding region. The coding region was that of the beta-tubulin paralogs 56D, 60D, 65B, 85D, and 97EF with either exon 4B or 4C. As described above, *UASt-betaTub56D* expression resulted in complete rescue of *betaTub56D*^*NP0949*^/*Df(2R)BSC782* first instar larval lethality (Fig. 5A). In contrast, *UASt-betaTub97EF(4B)* and *UASt-betaTub97EF(4C)* provided only limited rescue. Lethality still occurred, but during the second larval instar rather than already during the first (Fig. 5A). While such a partial rescue was also obtained with *UASt-betaTub65B*, the others (*UASt-betaTub60D* and *UASt-betaTub85D*) did not appear to cause rescue (Fig. 5A).

Additional experiments with the *UASt* transgenes revealed that ectopic expression of some beta-tubulin paralogs is highly toxic in an otherwise normal background. When the *UASt* transgenes were expressed with *ey-GAL4* during eye development for example (Fig. 5B), the beta-Tubulin 97EF isoforms were more toxic than all the other beta-tubulin paralogs. While the variant with the 4C exon was lethal, the 4B version resulted in small deformed and rough eyes, indicating that the two beta-Tubulin 97EF isoforms are functionally distinct.

When *mat-4-GAL-VP16* was used to drive *UASp-betaTub97EF(4B)* during oogenesis, female sterility was observed and the dumping of nurse cell cytoplasm into the oocyte that normally occurs during stage 10B was inhibited (Fig. 5C,D). Such a dumpless egg chamber phenotype was not obtained with *UASp-betaTub85D* or *UASp-betaTub60D* (Fig. 5D). Immunolabeling confirmed that ectopic expression of both the toxic beta-Tubulin 97EF and the non-toxic beta-Tubulin 60D occurred as expected in the germline starting during stage 2 (Fig. S5). During wild-type oogenesis, both beta-Tubulin 97EF and beta-Tubulin 60D are normally expressed in somatic follicle cells but not in the germline (Fig. S5).

The C-terminal tails of tubulin paralogs are the most divergent regions and different types of posttranslational modifications occur predominantly in this region, potentially generating a tubulin code that might direct interactions with specific sets of MAPs and motor proteins. To evaluate whether the functional difference between beta-Tubulin 97EF(4B) and beta-Tubulin 56D reflects sequence changes within the core or the C-terminal tail, we performed *betaTub56D* mutant rescue experiments with *UASt* transgene encoding tail-swopped variants. This assay indicated that the functional difference between beta-Tubulin 97EF(4B) and beta-Tubulin 56D reflects alterations within both the core and the C-terminal tail regions (Fig. 5A). We conclude that *betaTub97EF* is not just expressed in a specific pattern but that it also codes for a beta-tubulin with unique functions as a result of sequence changes within both the tubulin core and the C-terminal tail region.

For further analysis of the functional differences between beta-Tubulin 97EF and beta-Tubulin 56D, we studied larval hemocytes. Since S2R+ cells have an expression profile most similar to hemocytes (Cherbas et al., 2011), it appeared likely that larval hemocytes express *betaTub97EF* in a temperature-regulated manner. Indeed, immunolabeling confirmed that larval hemocytes do not only express *betaTub56D* but also *betaTub97EF* (Figs 2D, 6A). Moreover, the beta-Tubulin 97EF level was significantly increased at 14°C compared to 29°C (Fig. 6B). By exploiting larval hemocytes from *betaTub56D* mutants in combination with expression of *UASt* tubulin transgenes, we were able to alter the relative abundance of beta-Tubulin 97EF and beta-Tubulin 56D even at the optimal temperature of 25°C (Fig. 6A). Moreover, coexpression of *UASp-EB1-GFP* allowed quantitative analyses of some aspects of microtubule dynamics *in vivo*. EB1-GFP accumulates on the plus ends of growing microtubules (Mimori-Kiyosue et al., 2000). Therefore, we used hemocytes from second instar larvae with mutations in *betaTub56D* (*betaTub56D*^*NP0949*^/*Df(2R)BSC782)* and *alphaTub84BP-GAL4* driven expression of *UASt-betaTub97EF(4B)* or *UASt-betaTub56D* and also *UASp-EB1-GFP* in some of the experiments. In the following, these hemocytes will be designated as *R-97EF* and *R-56D* (rescued by the 97EF and 56D paralog, respectively). In addition, we also analyzed hemocytes from siblings (*S-97EF* and *S-56D*, respectively) which expressed the *UAS* transgenes in balanced *Df(2R)BSC782* larvae with a functional endogenous *betaTub56D* copy on the balancer.

**Fig. 6.**
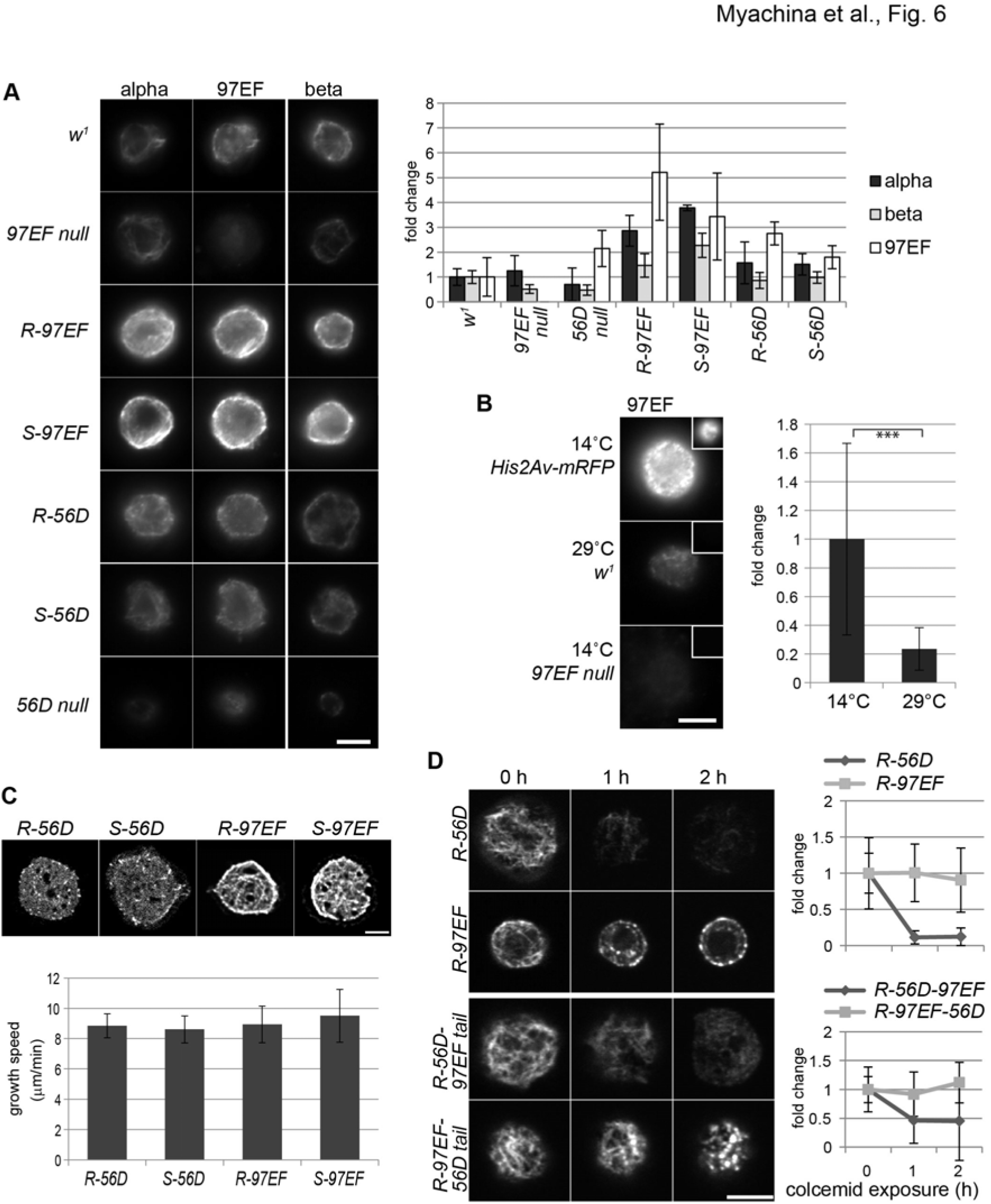
beta-Tubulin 97EF stabilizes microtubules in larval hemocytes. (A-D) Hemocytes were isolated from larvae with the indicated genotypes (97EF null = *betaTub97EF*^*MiMIC*^; 56D null = *betaTub56D*^*NP0949*^/*Df(2R)BSC782*; R-97EF and S-97EF = 56D null and *Df(2R)BSC782/*balancer, respectively, with *alphaTub84BP>betaTub97EF*; *R-56D* and S-56D = 56D null and *Df(2R)BSC782/*balancer, respectively, with *alphaTub84BP>betaTub56D*). (A) Hemocytes were double labeled with anti-beta-Tubulin 97E and anti-alpha-tubulin (left and middle columns) or only with anti-beta-tubulin (E7) (right column). Bar diagram represents average signal intensity (+/- s.d., n = 11). Intensities obtained in *w*^*1*^ were set to 1. (B) Hemocytes from larvae with the indicated genotypes after development at the indicated temperatures were released onto the same cover slip to enforce identical fixation and staining conditions. Nuclear His2Av-mRFP fluorescence (insets) was used for hemocyte classification. Bar diagrams indicate average background corrected anti-beta-Tubulin 97EF signal intensities (+/- s.d., n = 20). Signals observed at 14°C in *w*^*1*^ were set to 1. *** p < = 0.001 (*t* test). (C) Representative still frames after time lapse imaging of hemocytes expressing EB1-GFP. Bar diagram indicates average speed of EB1-GFP comet movements (+/- s.d., n = 11). (D) Hemocytes of indicated genotypes were treated with colcemid for the indicated time before fixation and staining with anti-alpha-tubulin. Graph represents average signal intensity (+/- s.d., n = 6). Signals observed at time 0 hours were set to 1 for each genotype. Scale bars 5 μm.

After immunolabeling, we quantified the signals obtained with anti-beta-Tubulin 97EF, DM1A (anti-alpha-tubulin) and E7 (anti-beta-tubulin) in the hemocytes (Fig. 6A). We point out that these signals primarily reflect the polymerized tubulin pool rather than the total cellular pool. Fixation seems to result in a loss of soluble tubulin, as suggested by the comparison of anti-tubulin immunolabeling signal intensities after brief pretreatment with either microtubule inhibitors (nocodazole, vinblastine) or buffer, respectively (Bossing et al., 2012; Schulman et al., 2013); see also Fig. 7). In addition, while non-specific background intensity resulting with anti-beta-Tubulin 97EF, as evident in *betaTub97EF* null mutant hemocytes, was subtracted from signal intensities obtained in the other genotypes, the extent of non-specific background resulting with DM1A (anti-alpha-tubulin) and E7 (anti-beta-tubulin) could not be assessed and subtracted. Moreover, non-linearity of detection might vary between different antibodies. Despite these limitations our signal quantification clearly revealed that expression of *UASt-betaTub97EF(4B)* and *UASt-betaTub56D*, respectively, has distinct consequences in larval hemocytes. *UASt-betaTub97EF(4B)* expression resulted in a pronounced increase of not only the signals obtained with anti-beta-Tubulin 97EF and E7, but also of those obtained with anti-alpha-tubulin (Fig. 6A, *R-97EF* and *S-97EF*). The anti-tubulin signals were particularly prominent in microtubule bundles along the cortex. In contrast, *UASt-betaTub56D* expression resulted in a more limited increase in signal intensities (Fig. 6A, *R-56D* and *S-56D*). Therefore, beta-Tubulin 97EF appears to stabilize microtubules. The posttranscriptional feedback preventing free tubulin heterodimer excess (Cleveland, 1988) likely contributes to the positive effect of *UASt-betaTub97EF(4B)* expression on anti-alpha-tubulin signal intensity.

**Fig. 7.**
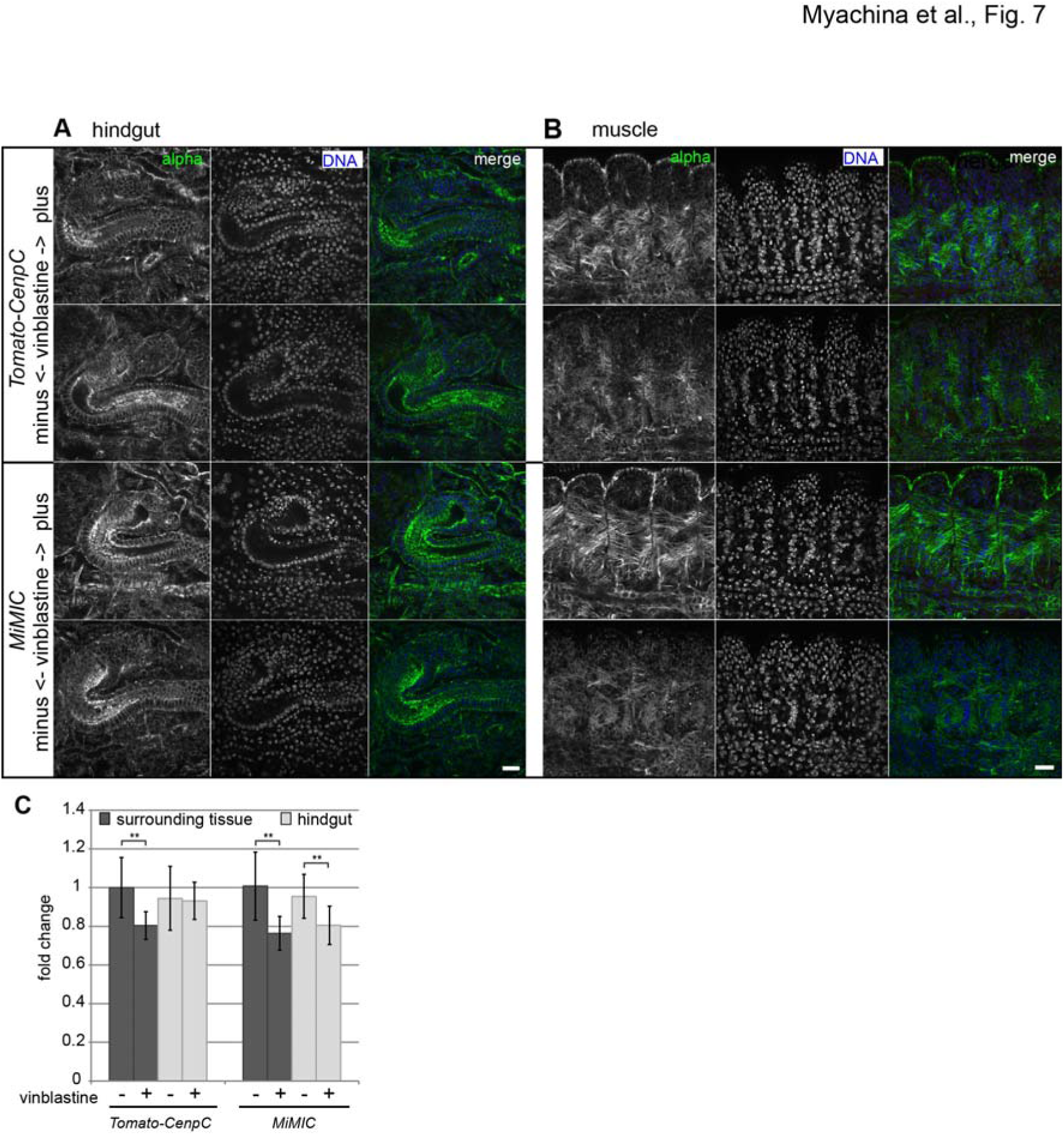
Increased microtubule stability in the embryonic hindgut depends on *betaTub97EF*. (A-C) Embryos (13-14 hours AED) with or without *betaTub97EF* gene function (*g-tdTomato-CenpC* and *MiMIC*, respectively) were exposed to vinblastine (+) or solvent only (-) before immunostaining with anti-alpha-tubulin. Sections illustrating the effect on anti-alpha-tubulin signal intensities in the hindgut and surrounding tissues (A) or in somatic muscles (B) are displayed. Scale bar 15 μm. (C) Bar diagram representing average anti-alpha-tubulin signal intensities in the hindgut and in the surrounding tissues (+/- s.d., n = 9). ** p < 0.01 (*t* test). Signals in controls were set to 1.

For additional comparison of the effects resulting from *UASt-betaTub97EF(4B)* and *UASt-betaTub56D* expression, we analyzed EB1-GFP comet movements. In *R-56D* and *S-56D* hemocytes, moving EB1-GFP comets were readily observed (Fig. 6C, Suppl Movie 1), similar as previously described in normal control hemocytes (Leśniewska et al., 2014). In contrast, EB1-GFP comets were far less evident in *R-97EF* and *S-97EF* hemocytes (Fig. 6C, Suppl Movie 2). In these cells, the mobile comets were partially masked by stably persisting EB1-GFP signals all along the microtubule network and most prominently at the cell cortex. Newly appearing EB1-GFP comets advanced mostly along the persisting bundles. Quantification of the dynamics of EB1-GFP comets did not reveal statistically significant differences in the effects of *UASt-betaTub97EF(4B)* and *UASt-betaTub56D* expression on the speed of EB1-GFP comet migration (Fig. 6C). These findings suggested that beta-Tubulin 97EF increases the amount of microtubules, but not by stimulating the microtubule growth rate. Thus it presumably decreases the catastrophe frequency and/or it increases the rescue frequency.

To further confirm that increased levels of beta-Tubulin 97EF stabilize microtubules, we analyzed their colcemid sensitivity. Colcemid inhibits microtubule growth and depolymerizes microtubules at higher concentrations (Ravelli et al., 2004). Colcemid was found to have a more pronounced effect on hemocytes expressing *UASt-betaTub56D* compared to those expressing *UASt-betaTub97EF(4B)* (Fig. 6D). In *R-56D* hemocytes, anti-alpha-tubulin signals were drastically lower after one hour incubation in the presence of colcemid. In contrast, signals were still high in *R-97EF* hemocytes even after 2 hours of incubation although in a more patchy distribution. Analogous analyses after expression of beta-Tubulin 56D and 97EF with swopped C-terminal tails, indicated that the microtubule stabilizing effect is mediated by the beta-Tubulin 97EF core and not by its C-terminal tail (Fig. 6D). Our results support the notion that beta-Tubulin 97EF stabilizes microtubules.

The analysis of microtubules in hemocytes indicated that microtubules containing beta-Tubulin 97EF are more stable than those built primarily from the major embryonic maternally provided beta-Tubulin 56D. Accordingly, the hindgut microtubules of *betaTub97EF* mutants (containing only maternally derived beta-Tubulin 56D) are predicted to be less stable than those in wild-type control embryos (containing beta-Tubulin 56D and beta-Tubulin 97EF). To address this prediction, we performed microtubule depolymerization experiments with embryos with or without *betaTub97EF* function. For these experiments, we used vinblastine because it depolymerized microtubules in embryos more effectively than colcemid (data not shown). To enforce equal conditions during vinblastine treatment as well as during subsequent fixation and immunolabeling, embryos with and without *betaTub97EF* function were combined into a pool before treatment onset. To identify the different genotypes during the microscopic analysis eventually, the embryos with *betaTub97EF* gene function had the *g-tdTomato-CenpC* marker gene, resulting in red fluorescent centromere signals. In contrast, the *betaTub97EF* mutants did not have this marker transgene. After vinblastine or mock treatment, embryos were stained with anti-alpha-tubulin and signal intensities were quantified. The quantification revealed that in the hindgut, vinblastine caused a significant reduction in the level of anti-alpha-tubulin signals but only in embryos lacking *betaTub97EF* function (Fig. 7). In contrast, in the tissues surrounding the hindgut and in the body muscles, vinblastine reduced the level of anti-alpha-tubulin signals to a comparable extent in embryos with and without *betaTub97EF* function (Fig. 7). These results indicate that microtubules in normal hindgut, where *betaTub97EF* expression is high, are more stable than in tissues, where *betaTub97EF* is at most marginally expressed. Moreover, this higher microtubule stability in the hindgut depends on *betaTub97EF* expression, as it is lost in *betaTub97EF* mutants.

## Discussion

The molecular mechanisms allowing acclimation to ambient temperature fluctuations are poorly understood. Here we have identified *betaTub97EF* as a temperature responsive gene in *Drosophila melanogaster*. At low temperature, *betaTub97EF* expression is increased. However, at the optimal temperature, as well as at lower temperatures, *betaTub97EF* is not expressed ubiquitously but restricted in a range of tissues including gut epithelial cells and hemocytes. Mutations that eliminate *betaTub97EF* function have a surprisingly mild effect, presumably reflecting functional redundancy with other co-expressed beta-tubulin paralogs. But without beta-Tubulin 97EF, microtubules are less stable. Conversely, microtubules are stabilized by increased *betaTub97EF* expression. Moreover, within the well-tolerated temperature range, embryogenesis of *betaTub97EF* mutants is more cold-than heat-sensitive. Therefore, upregulation of *betaTub97EF* at low temperature might contribute to acclimation by stabilizing microtubules.

A detailed understanding of the *betaTub97EF* contribution to the success of embryogenesis specifically at the lower end of the temperature range compatible with completion of the entire life cycle will require further work. Lack of beta-Tubulin 97EF in the gut epithelia might impair yolk utilization during late embryogenesis at low temperature. It might also affect hemocyte functions (Wood and Martin, 2017). Hemocytes are important for innate immunity, which is unlikely to be relevant for successful embryogenesis. However, they also provide functions essential for embryogenesis. They scavenge apoptotic corpses resulting from the abundant developmentally programmed cell death. Moreover, they are crucial for secretion and remodeling of much of the extracellular matrix in embryos.

Microtubules are ubiquitous and temperature change will affect them in all cells. Irrespective of temperature change, microtubule properties are carefully controlled in animal cells, not only by alterations in tubulin isoform composition, but also by myriads of interacting factors. Thereby their characteristics in different cell types and intercellular subsets are modified to serve a wide range of functions. Acclimation of these diverse microtubule systems to temperature change is therefore likely demanding and impossible to achieve simply and exclusively by global upregulation of a single tubulin paralog. Presumably, tissue-specific upregulation of *betaTub97EF* is one of many mechanisms for microtubule acclimation in *D. melanogaster*.

Beyond acclimation to low temperature, *betaTub97EF* expression occurs also at optimal and elevated growth temperatures. *betaTub97EF* is one of the five beta-tubulin paralogs present in the genome of *D. melanogaster*, which includes five alpha-tubulin paralogs. The human genome hosts almost twice as many alpha- and beta-tubulin genes (9 and 8, respectively) and genetic analysis during the last 10 years have uncovered an increasing number of disease-associated mutations primarily in the context of neurological disease and cancer (Bahi-Buisson et al., 2014; Ludueña and Banerjee, 2008; Wang et al., 2017). Characterization of these genes and associated phenotypes, also in mouse models, has fully confirmed the conclusions from pioneering analyses of tubulin paralog functions in *D. melanogaster*. Tubulin paralogs have distinct but often partially overlapping patterns of expression and functions. The understanding of how amino acid sequence alterations affect tubulin function has progressed slowly. Production of pure recombinant tubulin isoforms in quantities sufficient for studies of microtubule polymerization and dynamics *in vitro* has been achieved only very recently (Pamula et al., 2016; Sirajuddin et al., 2014; Ti et al., 2016; Valenstein and Roll-Mecak, 2016; Vemu et al., 2016; Widlund et al., 2012). As the microtubule interactome in cells is highly complex (Hughes et al., 2008; Yu et al., 2016) phenotypic analyses *in vivo* will remain important as well.

Here we have used a series of transgenes for *GAL4*-regulated expression of all the *D. melanogaster* beta-tubulin paralogs. As the transgenes are identical except for the coding sequences, differential effects most likely reflect functional specialization at the protein level. Expression with *ey-GAL4* (and additional drivers including *en-GAL4* and *da-GAL4*, data not shown) indicated that excess *betaTub97EF* is more toxic than that of the other beta-tubulin paralogs. Moreover, *betaTub97EF-4C* is more detrimental than *betaTub97EF-4B*. These two *betaTub97EF* isoforms, resulting from mutually exclusive splicing of the fourth exon, differ at 13 positions within an internal segment of 40 amino acids, but their N- and C-terminal regions are identical. Sequence divergence between tubulin paralogs is generally most extensive within the tails, in particular C-terminally where also several distinct post-translational modifications occur (Gadadhar et al., 2017; Janke, 2014; Yu et al., 2015). Therefore, the C-terminal tails (CTTs) are thought to make important paralog-specific contributions, as fully confirmed recently (Sirajuddin et al., 2014). Our comparison of the 97EF and 56D beta-tubulins also revealed that CCTs contribute to paralog-specific function. While 56D with an 97EF-CTT behaved like normal 56D, 97EF with an 56D-CTT had a toxicity intermediate between 97EF and 56D, suggesting that the beta-Tubulin 97EF core and tail cooperate, as demonstrated most elegantly with the 56D and 85D paralogs (Popodi et al., 2008). Beyond CTT contributions, differences within the tubulin core were found to have more profound effects. The core is responsible for the characteristic microtubule stabilizing effect of beta-Tubulin 97EF in larval hemocytes. Similarly, a study with chimeric tail-swapped human beta-tubulins IIB and III concluded that differences between the cores were largely responsible for the different microtubule dynamics observed *in vitro* (Pamula et al., 2016).

Our results indicate that beta-Tubulin 97EF expression does not affect the microtubule growth rate but it results in microtubules that are less prone to destabilization by colcemid and vinblastine. As beta-Tubulin 97EF appears to be enriched in cortical microtubules, it might confer resistance against cortical factors that stimulate catastrophe. Cortical enrichment was apparent after double labeling of larval hemocytes with anti-alpha–tubulin and anti-beta-Tubulin 97EF. However, locally distinct epitope masking effects, for example by CTT-associated proteins or modifications, are clearly not excluded. Judging from the pattern of microtubules induced by beta-Tubulin 97EF overexpression in hemocytes, this tubulin might also facilitate microtubule bundling (Molodtsov et al., 2016). Interestingly, while ectopic *betaTub60D* expression in the germline did not affect oogenesis, *betaTub97EF* specifically blocked dumping of nurse cell cytoplasm into the oocyte during mid-oogenesis, i.e. long after the onset of ectopic expression. Dumping is critically dependent on actin and myosin. Microtubule inhibitors, which clearly inhibit some aspects of transport from nurse cells to oocyte, do not block the dumping process (Gutzeit, 1986; Lu et al., 2016; Roth and Lynch, 2009; Wheatley et al., 1995, Huelsmann, 2013 #7420). An excess of cortical microtubule bundles caused by ectopic beta-Tubulin 97EF might therefore inhibit dumping by interference with normal myosin-mediated contraction of the cortical actin network in nurse cells. Evidently, further analyses will be required to evaluate our speculative proposals. It is clear, however, that beta-Tubulin 97EF has interesting properties that are remarkably distinct from those of other *D. melanogaster* paralogs.

## Materials and methods

### *Drosophila* strains

Several stocks were obtained from the Bloomington Drosophila Stock Center (Indiana University, Bloomington, IN, USA) and the Kyoto Stock Center (Kyoto Institute of Technology, Kyoto, JP): *betaTub97EF*^*MiMIC*^ (#41519) (Venken et al., 2011), *betaTub97EF*^*MBO1877*^ (# 24452) and *betaTub97EF*^*MBO3812*^ (#24275) (Metaxakis et al., 2005), *betaTub56D*^*EY12300*^ (#20734) (Bellen et al., 2004); *Df(2R)BSC782* (#27354) and *Df(3R)BSC460* (#24964) (Cook et al., 2012), *betaTub56D*^*NP0949*^ (#103830) (Hayashi et al., 2002), *betaTub97EF*^*EGFP*^ was generated as described in further detail (Supplementary Information). Two strains with *UASt-Tub xy* transgenes were obtained from FlyORF (University of Zurich, Zurich, Switzerland): *M{UAS-betaTub85D.ORF}ZH-86Fb* (F001298) and *M{UAS-betaTub97EF(4B).ORF}ZH-86Fb* (F001587) (Bischof et al., 2013). Additional *UASt-Tub* lines (*M{UAS-betaTub56D.ORF}ZH-86Fb, M{UAS-betaTub60D.ORF}ZH-86Fb, M{UAS-betaTub65B.ORF}ZH-86Fb, M{UAS-betaTub97EF(4C).ORF}ZH-86Fb, M{UAS-betaTub56D-97EFtail.ORF}ZH-86Fb M{UAS-betaTub97EF(4B)-56Dtail.ORF}ZH-86Fb*) as well as the *UASp-Tub* lines *M{UASp-betaTub97EF(4B).ORF}ZH-86Fb* and *M{UASp-betaTub60D.ORF}ZH-86Fb* were generated as described in further detail (Supplementary Information). *UASp-EB1-GFP* (Jankovics and Brunner, 2006) was kindly provided by D. Brunner (University of Zurich, Zurich, Switzerland). The GAL4 driver and marker lines have been described previously: *ey-GAL4* (*5/8)* (Hazelett et al., 1998), *matα4-GAL-VP16(V2H)* (Hacker and Perrimon, 1998), *alphaTub84BP-GAL4 (LL7)* (#5138) (Lee and Luo, 1999) and *gi2xtdTomato-Cenp-C* II.3 (Althoff et al., 2012).

For analysis of the extent of *betaTub56D* mutant development supported by other beta-tubulin paralogs, the following crosses were set up. Virgins *Df(2R)BSC782/CyO, Dfd-YFP; UASt-betaTubXY* with XY = 56D, 60D, 65B, 85D, 97EF(4C) or 97EF(4C) were crossed with males w*; *betaTub56D*^*NP0949*^/*CyO, Dfd-GMR-nvYFP*; *alphaTub84BP-GAL4*/*TM6B, Hu, Dfd-YFP*. Eggs were collected from these crosses on either apple agar plates or in tubes with standard fly food. Collection plates were inspected at intervals under a fluorescence stereomicroscope allowing identification of larvae without the *Dfd-YFP* balancer chromosomes.

As control strain, *w* was used. *betaTub97EF*^*MiMIC*^ was backcrossed for four generations to *w* before comparison of the temperature sensitivity of development.

The rate of larval hatching from eggs was determined as described previously (Radermacher et al., 2014). Eggs were collected during 60 minutes on apple agar plates at 25°C and aged at this temperature for 5 hours. The collection plates with the eggs were then divided into sectors. The eggs on one of the sectors were fixed immediately with methanol and stained with a DNA stain to determine the fraction of unfertilized eggs. The other sectors were shifted to different temperatures and the fraction of hatched and unhatched eggs was determined eventually with the help of a stereomicroscope. The hatch rate was calculated as the ratio of the number of hatched eggs to the total number of eggs deposited on the sector minus the number of unfertilized eggs. For the analysis of the rate of development to the adult stage, eggs were also collected during 60 minutes at 25°C followed by ageing at this temperature for 5 hours. Sectors of the collection plates with eggs that had been counted were then separated into bottles containing standard fly food and shifted to different temperatures. The number of eclosed flies was determined eventually. The eclosion rate was calculated as a ratio between the number of eclosed flies to the total number of eggs minus the number of unfertilized embryos.

### Transcript analyses

The analysis of temperature effects on the transcriptome of cultured cells will be described in detail elsewhere. Data shown here was obtained with S2R+ cells which were cultured as described previously (Radermacher et al., 2014). Aliquots were shifted 24 hours after plating at 25°C to different temperatures (11, 14, 25 and 30°C). 24 hours later RNA was extracted, followed by probe synthesis and microarray hybridization (*Drosophila* gene expression 4x44K microarrays, G2519F-021791, Agilent Technologies). Three independent replicas were analyzed and after normalization of signal intensities across replicates average hybridization signals were determined. In contrast to all other tubulin paralogs, signals for the testis-specific *betaTub85D* paralog were too low to be included in the S2R+ transcriptome analysis (Fig. 1A).

For qRT-PCR analysis, S2R+ cells were treated in the same way as for the microarray experiments. For qRT-PCR analysis with *w*^*1*^ embryos, eggs were collected at 25°C for 3 hours. After additional ageing at 25°C for 3 hours, they were shifted to 14, 25 and 30°C for 40, 10 or 8 hours, respectively. As a result, embryos were all at stage 16 of embryogenesis irrespective of the applied temperature regime. An aliquot of each sample was fixed for controlling the developmental stage after DNA staining by microscopy. Further details concerning the qRT-PCR analyses, including primer sequences, are described in Supplementary Information.

### Antibodies

Detailed information on all antibodies and dilutions applied during immunoblotting and immunolabeling is given in the Supplementary Information. The rabbit antibodies against beta-Tubulin 97EF were generated with a synthetic peptide corresponding to last 17 amino acids (AEQEGYESEVLQNGNGE) coupled to KHL (Moravian Biotechnology, Brno, Czech Republic). Antibodies were affinity purified. Rabbit antibodies against beta-Tubulin 60D (Leiss et al., 1988) were kindly provided by R. Renkawitz-Pohl (University of Marburg, Marburg, Germany).

### Immunoblotting

Total extracts from cells, embryos, larvae and adults were prepared as described in further detail in the Supplementary Information and resolved by gel electrophoresis. Embryos and larvae aged at different temperatures were processed with solutions that had been pre-equilibrated to the temperatures used for aging. For probing samples with different anti-tubulin antibodies, separate gels and immunoblots were used for each antibody in order to avoid potential masking effects that might arise during serial probing of a single immunoblot. Immunoblot signals were developed using chemiluminescence (Amersham ECL, GE Health Care Life Sciences) and captured using a CCD camera. For quantification of immunoblot signals a dilution series of a reference extract was resolved along with the test samples. Quantification of signals by densitometry was done with ImageJ as described previously (Radermacher et al., 2014).

### Microscopic analyses

Embryos were immunolabeled as described before (Radermacher et al., 2014). For vinblastine treatment, *gi2xtdTomato-Cenp-C* II.3 and *betaTub97EF*^*MiMIC*^ embryos were collected and aged to 11-13 hours AED. Pooled embryos were dechorionated with 3% sodium hypochlorite, washed extensively with 0.7% NaCl, 0.07% Triton X-100 and transferred into an Eppendorf tube. 1.5 ml of a 1:1 mixture of heptane and Schneider’s medium were added. Moreover, 3 μl of a vinblastine (Sigma, V1377) stock solution (5 mM in methanol) or 3 μl methanol in case of control aliquots were added. The Eppendorf tubes were slowly rotated end over end for 20 minutes at room temperature. Thereafter, embryos were fixed with methanol.

Preparation and immunolabeling of organs from larvae and adults was performed using standard procedures as described in further detail in Supplementary Information.

Hemocytes were isolated as previously described (Sampson and Williams, 2012) using second instar larvae (*R-56D, S-56D, R-97EF*, *S-97EF*) or first instar larvae (*betaTub56D*^*NP0949*^/*Df(2R)BSC782*). The larvae were collected in a drop of 150 μl Schneider’s medium on a glass slide that had been pre-coated with 0.5 mg/ml concanavaline A (Sigma, C7275) in phosphate buffer (9 mM NaH_2_PO_4_, 1 mM Na_2_HPO_4_, 1 mM CaCl2, pH 6.0). Larval cuticles were gently ruptured with forceps. This allowed migration of hemocytes out of the larvae and their subsequent initial attachment to the cover slip. After 15 minutes at room temperature, larvae were discarded. Before fixation or live imaging, hemocytes were allowed to spread during a 60 minute incubation at 25°C. In case of colcemid treatment, the compound was added to final concentration 10 μM in Schneider’s medium. Hemocytes were fixed either with 4% paraformaldehyde in phosphate buffered saline (PBS) at room temperature or (for Fig. 6D) with methanol containing 1 mM EGTA for 10 minutes at −20°C (Hoogenraad et al., 2000).

Images from fixed samples were acquired with an Olympus FluoView 1000 laser-scanning confocal microscope with a 60x/1.35 objective (Figs 2C, 3D, 4C,D, 6D,E, 7A,B), with a 40x/1.3 objective (Sup. Figs. 3B,C, 5) or with a Zeiss Cell Observer HS wide-field microscope with a 63x/1.4 objective (Figs 2B,D, 6A,B), or with 16x/0.5 objective (Fig. 5C). Z-stacks were acquired. Single sections are presented in all images, except in Fig. 3A, where maximum intensity projections of relevant regions made with Imaris (Bitplane) are displayed. The intensities of anti-tubulin signals in hindgut in Fig. 4C-E were measured with ImageJ in maximum projections of 5 slices taken within the middle of the hindgut tube. The intensities of anti-alpha-tubulin staining in hindgut and other embryonic tissues (Fig. 7) were measured in single slices with ImageJ after manual selection of the regions of interest. Image z-stacks of eyes (Fig. 5B) were acquired with a Zeiss AxioCam HRc camera and converted into extended focus images using Helicon Focus Software (Helicon Soft Ltd.).

Time lapse imaging of EB1-GFP comets in live hemocytes was done with a VisiScope spinning disc confocal microscope (Visitron) on an Olympus IX83 microscope stand with a 100x/1.4 objective. Three sections with 0.1 μm spacing were acquired at 1 sec intervals. Signals in case of *R-97EF* and *S-97EF* were deconvolved using Huygens Software (Scientific Volume Imaging b.v.). Quantification of microtubule dynamics after sum of intensity projections was done with u-track software (Jaqaman et al., 2008). To differentiate the moving EB1-GFP comet signals from the stationary EB1-GFP signals characteristically present in the *R-97EF* and *S-97EF* hemocytes along microtubule bundles, image sequences were processed arithmetically in ImageJ as follows. In a first variant image sequence, the first four frames were deleted. The last four frames were deleted in a second variant. By subtracting variant 2 from variant 1, we arrived at an image sequence from which background was subtracted (Image J, rolling ball radius 2 pixels) before analysis with u-track.

Characterization of the hindgut ultrastructure in *w*^*1*^ and *betaTub97EF*^*MiMIC*^ after development at 25°C to 15-16 hours AED by transmission electron microscopy and FIBSEM was achieved as described in detail in Supplementary information.

## Acknowledgements

We thank Renate Renkawitz-Pohl for anti-beta-Tubulin 60D and Nikola Linsi for his contributions to antibody characterization. We are also grateful to Rita Lecca and the Functional Genomics Center Zurich for supporting microarray hybridizations, to Martin Moser for qRT-PCR support, as well as to Andres Käch and the Center for Microscopy and Image Analysis (University of Zurich).

## Competing interests

No competing interests declared.

## Author contributions

F.M. performed all experiments (except microarray analyses and part of electron microscopy), analyzed data, designed experiments together with C.F.L. and contributed to manuscript writing. F.B. performed expression profiling. J.B. supported the generation of transgenic strains. M.K. performed electron microscopy, C.F.L conceived and supervised the study, analyzed data, and wrote the manuscript.

## Funding

This work was supported by the Swiss National Science Foundation (grant 31003A_152667 to C.F.L.).

---

**Movie S1. EB1-GFP comets in hemocytes expressing different beta-Tubulin isoforms.** Larval hemocytes (designated as *R-56D, S-56D, R-97EF, and S-97EF* as in the main text) were isolated from *betaTub56D*^*NP0949*^/*Df(2R)BSC782* (*R*) or balancer/*Df(2R)BSC782* (*S*) larvae with *alphaTub84BP-GAL4* driven expression of *UASp-EB1-GFP* and either *UASt-betaTub56D* or *UASt-betaTub97EF*. Sum intensity projections of three optical sections acquired at 1 second intervals are shown.

## Supplementary Materials and Methods

### Plasmids

Constructs for bacterial expression of GST fusion proteins with N- and C-terminal extensions corresponding to the terminal sequences of the different *Drosophila* beta-tubulin paralogs were made using the pET-21d vector (Novagen). Gene blocks (Integrated DNA Technologies) coding for the N- and C-terminal regions as indicated in Fig. S1 were inserted. The sequences of these gene blocks are given in Table S1. The gene blocks carried an NcoI and a NotI site at the 5’ and 3’ end, respectively. These sites were used for cloning into the corresponding sites of pET-21d. Moreover, a linker sequence with a HindIII and BamHI site was present in the gene blocks in between the N- and C-terminal coding sequences. These sites were used for the insertion of the GST coding region that was amplified with CL204 and CL205 from the pGEX-2T vector (Smith and Johnson, 1988).

**Fig. S1.**
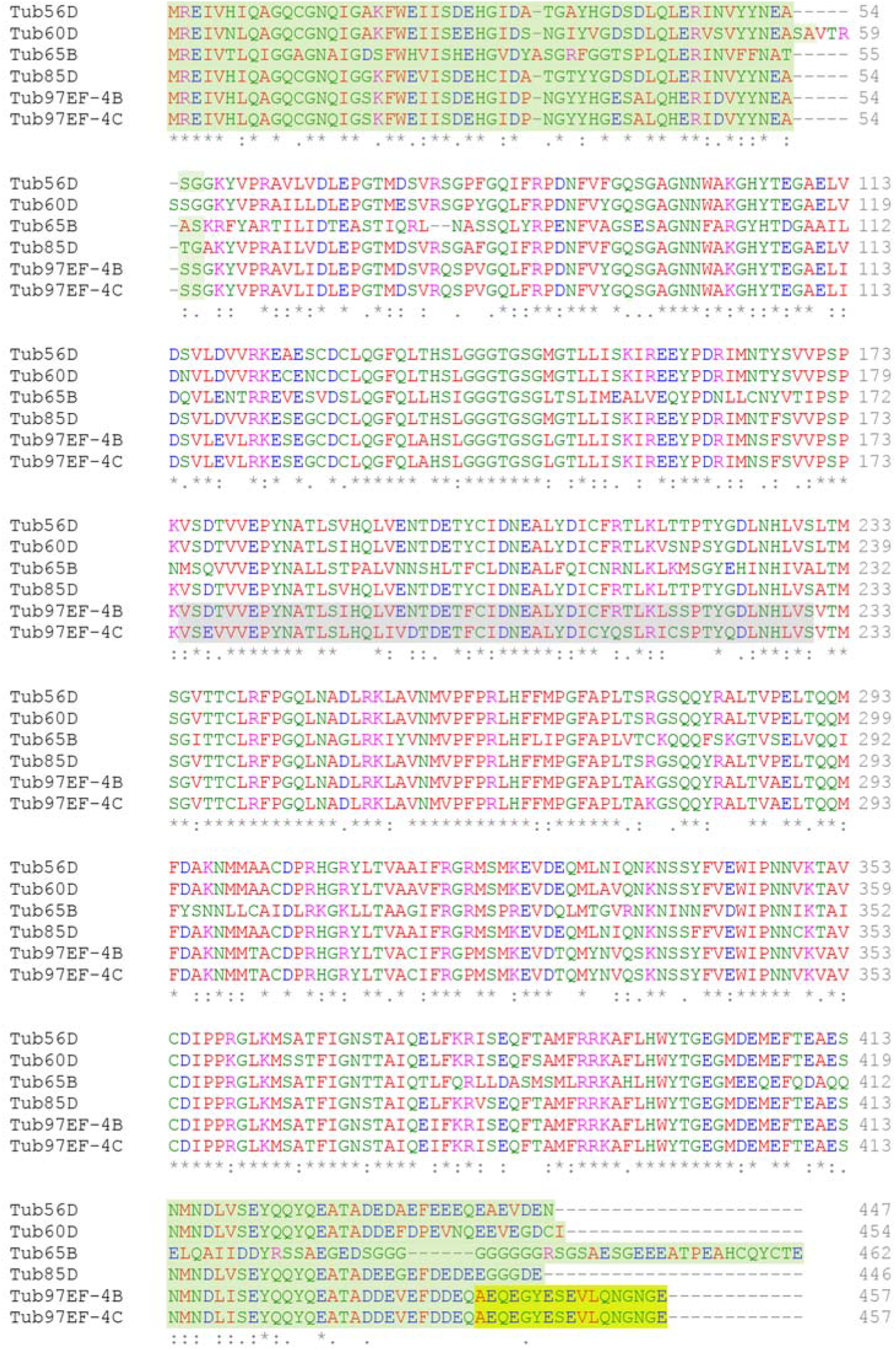
Comparison of amino acid sequences of *Drosophila* beta-tubulin paralogs. Highlighted in yellow is the sequence of the peptide used for the production of a beta-Tubulin 97EF-specific antibody. The region encoded by the mutually exclusive exons 4B and 4C of beta-Tubulin 97EF are indicated by grey shading. The N- and C-terminal sequences (green shading) are less conserved than the core domains. These terminal sequences were present in the GST fusions which were expressed in bacteria for the characterization of various antibodies (see Fig. S2).

**Fig. S2.**
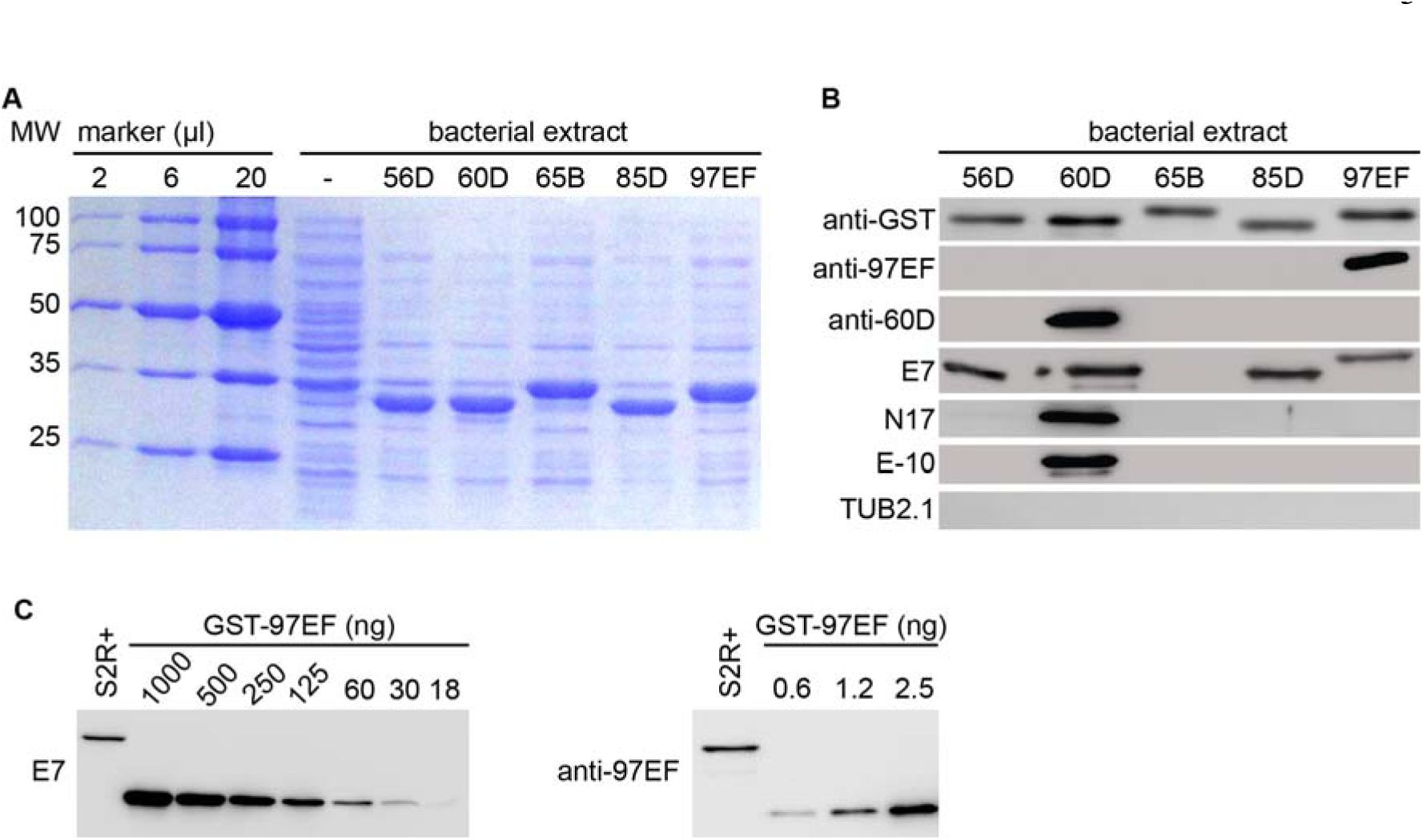
Characterization of various anti-beta-tubulin antibodies. (A) Bacterial extracts containing GST fusion proteins with distinct N- and C-terminal extensions corresponding to the N and C-terminal tails of the indicated *Drosophila* beta-tubulin paralogs were resolved by SDS-PAGE and analyzed by Coomassie Blue staining. In addition, a control extract (-) from bacteria without an expression construct, as well as a dilution series with known amounts of molecular weight marker proteins was analyzed. (B) Extracts with GST beta-tubulin tail fusions (as in A) were analyzed by immunoblotting with the indicated antibodies. 100 ng of GST fusion protein per lane was loaded in case of the immunoblots with anti-GST, E7, E-10, and dN-17, while 5 ng was loaded for the analyses with anti-97EF and anti-60D. (C) Total extracts of S2R+ cells grown at 25°C were analyzed by immunoblotting with anti-beta-Tubulin 97EF and anti-beta-Tubulin E7. 20 μg of total cellular protein was loaded in case of the anti-97EF immunoblot and 10 μg in case of the E7 immunoblot. For determination of the amount of beta-Tubulin 97EF and of total beta-tubulin in S2R+ cells a dilution series of the bacterial extract containing a known amount of the GST fusion protein with beta-tubulin 97EF tails was resolved and analyzed on the same immunoblots.

**Fig. S3.**
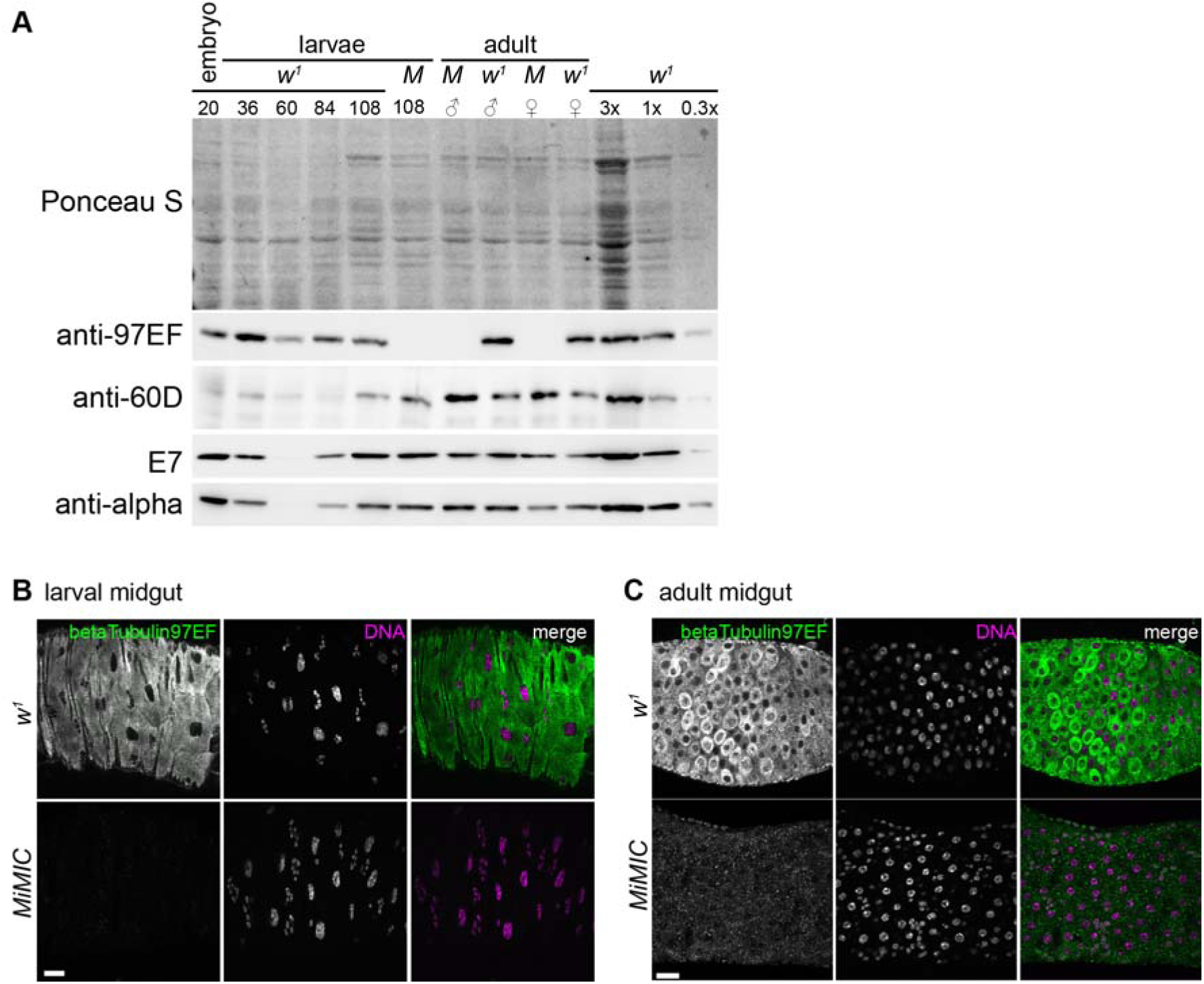
*betaTub97EF* expression in larval and adult tissues. (A) Total extracts prepared from either late embryos (embryo), larvae or adults were resolved by SDS-PAGE and analyzed after Western blotting by Ponceau S staining and probing with the indicated antibodies. Samples were from either the *w*^1^ control strain or *betaTub97EF*^*MiMIC*^ null mutants (M for *MiMIC*). Embryonic and larval extracts were prepared from samples aged for the indicated time (hours after AED). 36, 60, 84 and 108 hours AED correspond to mid-first, mid-second, early and late third instar, respectively. In case of adults, flies (0-1 day after eclosion) were separated according to sex before extract preparation. Equal amounts of protein were loaded in all lanes except for the last three where a dilution series of the *w*^*1*^ larval extract (108 hours AED) was loaded as internal reference for quantification. (B,C) Immunostaining of the gut from *w*^*1*^ and *betaTub97EF*^*MiMIC*^ animals during third larval instar wandering stage (B) or from adults (C) with anti-beta-Tubulin 97EF and a DNA stain. Scale bar 20 μm.

**Fig. S4.**
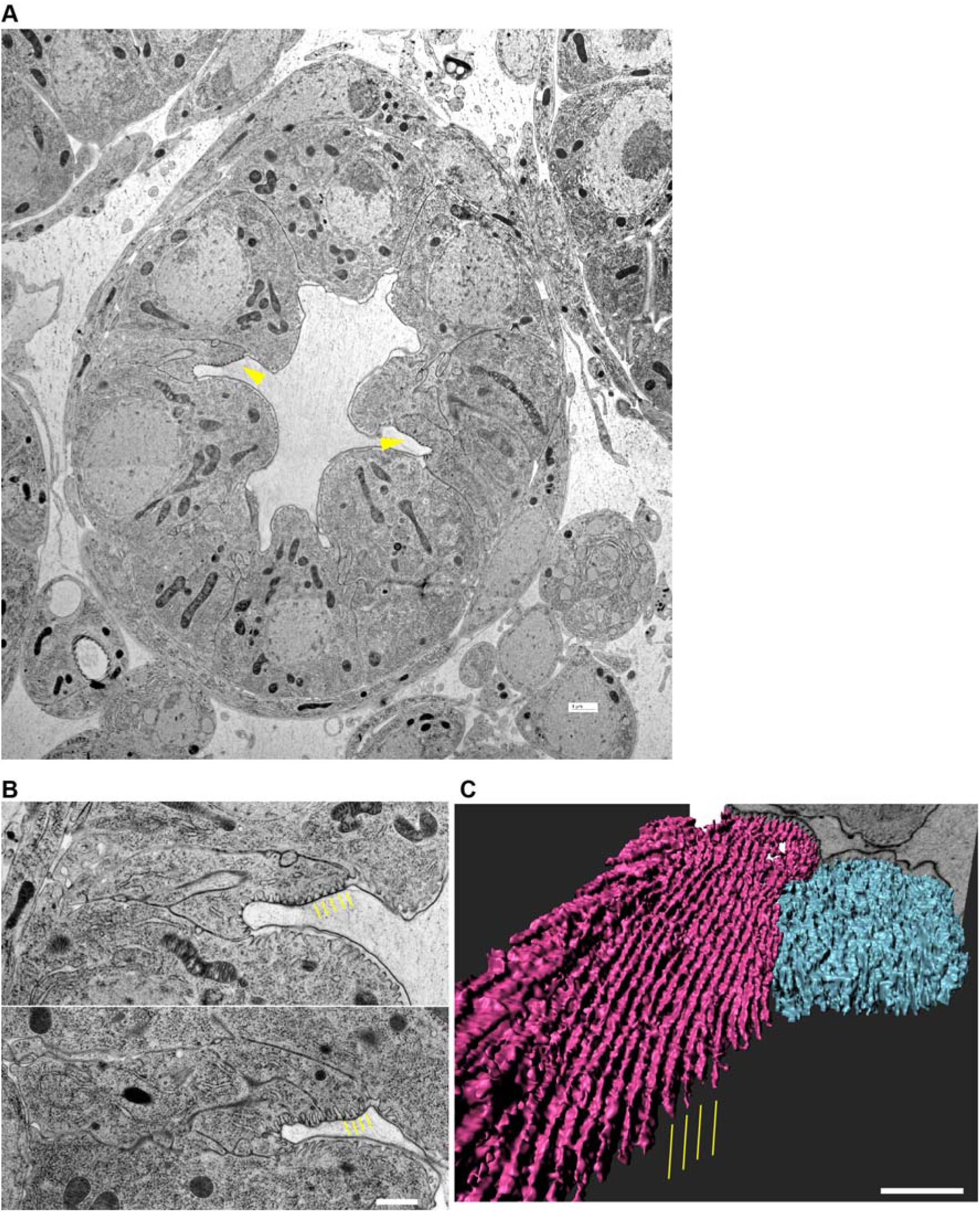
Boundary cells in the embryonic hindgut form longitudinally extended undulae. (A) Embryos at stage 16 were fixed by high-pressure freezing and processed for TEM and FIBSEM analysis. The cross section through a control hindgut reveals the characteristic apical structures in boundary cells (yellow arrowheads). Scale bar 1 μm. (B) The boundary cells in both control (top) and *betaTub97EF*^*MiMIC*^ mutants (bottom) differentiate the characteristic apical structures (yellow dashes). Scale bar 1 μm. (C) 3D-reconstruction of the apical surface of a boundary cell (purple) and a neighboring principal cell (cyan) after FIBSEM reveals that the characteristic apical structures of boundary cells represent highly organized longitudinal undulae, which are absent from principal cells (cyan). Scale bar 0.5 μm.

**Fig. S5.**
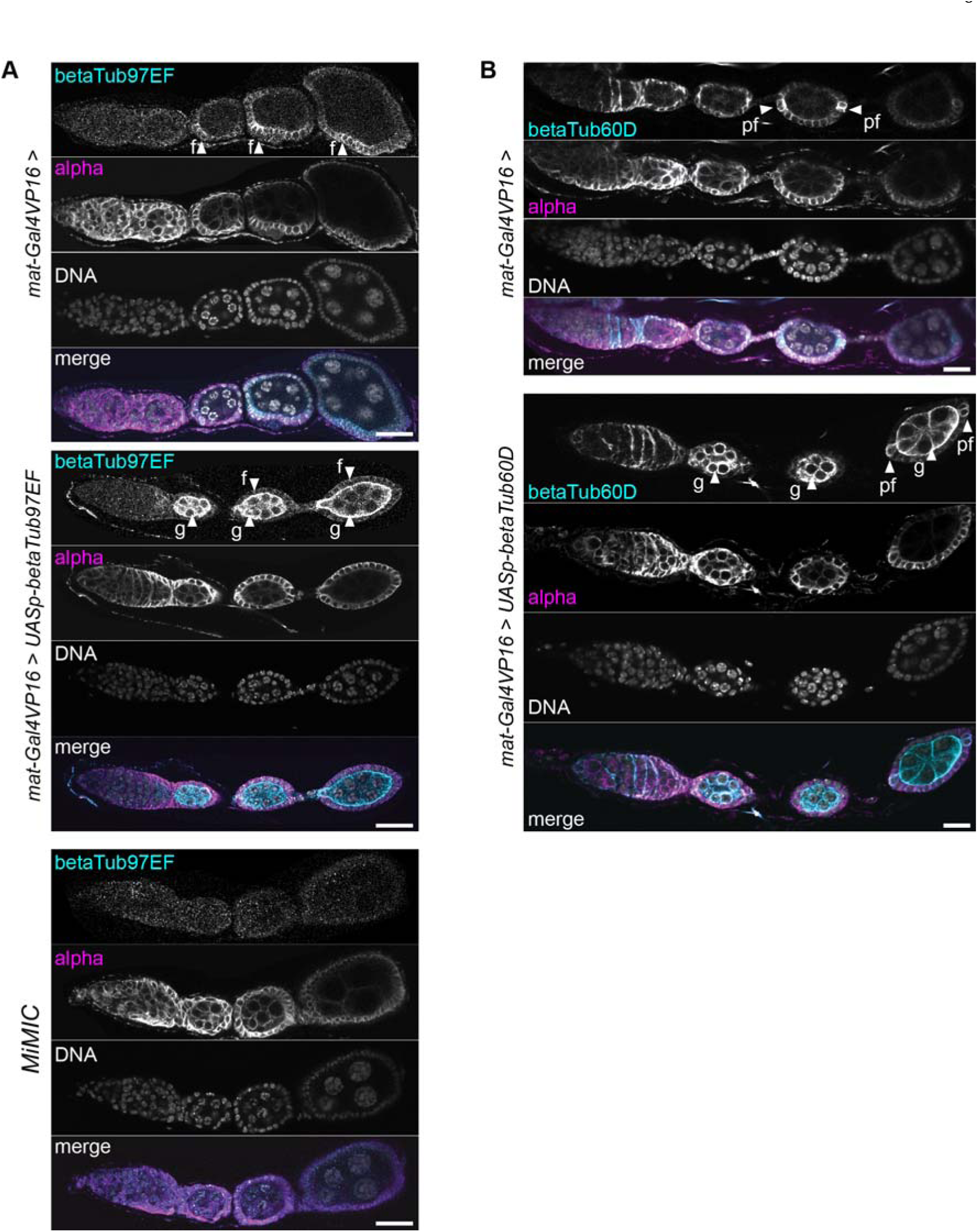
Ectopic *betaTub97EF* and *betaTub60D* expression in the female germline during oogenesis. Ovaries were isolated from females with the indicated genotypes (MiMIC = *betaTub97EF*^*MiMIC*^). After fixation, DNA staining and immunolabeling with indicated antibodies were performed. The early stages of oogenesis in representative ovarioles are shown. *mat-GAL4VP16* drives expression of *UASp* transgenes in the germline, starting during stage 2 of oogenesis. Arrowheads indicate expression in follicle cells (f), germline cells (g) and polar follicle cells (pf). Scale bars 20 μm.

Additional *Drosophila* ORFeome constructs were made with the previously described cloning strategy (Bischof et al., 2013). The coding regions of *Drosophila* beta-tubulin paralogs were amplified from the following cDNA plasmids (Drosophila Genome Resource Center, Indiana University, Indiana, USA) with the indicated primer pairs (see Table S1 for sequences): *betaTub56D* from pOT2_LD43681 (Rubin et al., 2000) with FM200/FM201, *betaTub60D* from pFLC1_RE53159 (Stapleton et al., 2002) with FM116/FM117, *betaTub65B* from pOTB7_AT27896 (Stapleton et al., 2002) with FM122/FM123, *betaTub97EF-4C* from pOT2_Tub97EF_4C with FM113/FM114. pOT2_Tub97EF_4C was made using pOT2-LP10436 (Rubin, 2000 #7431), which contains a *betaTub97EF-4B* cDNA, by exchanging a BamHI and EheI fragment including the exon 4B with a gene block (Integrated DNA Technologies) containing exon 4C instead of 4B (see Table S1 for sequence).

For GAL4-regulated expression in the female germline, we generated transgene constructs using the vector pUASp-K10attB kindly provided by B. Suter (University of Berne, Switzerland). The coding regions of beta-tubulin paralogs were inserted into the SfiI and XbaI sites of this vector after enzymatic amplification from the following cDNA plasmids with the indicated primer pairs (see Table S1 for sequences): *betaTub60D* from pFLC-1-RE53159 with FM212/FM213; *betaTub85D* from pOT2-GH62051 with FM214/FM215, *betaTub97EF-4B* from pOT2-LP10436 with FM208/FM209.

### Generation of transgenic *Drosophila* strains

The additional *Drosophila* ORFeome constructs, as well as the pUASp-K10attB constructs described above were injected into *y^1^ M{vas-int.Dm}ZH-2A w^*^; M{3xP3-RFP.attP}ZH-86Fb* (Bischof et al., 2007). F1 progeny was crossed to *yw* and progeny was screened for *w^+^* flies, followed by establishment of transgenic lines.

The reporter line *betaTub97EF*^*EGFP*^ was generated by recombinase mediated cassette exchange using *betaTub97EF*^*MiMIC[MI06334]*^ (Venken et al., 2011). The cassette present in Mi{MIC}[MI06334] was replaced with the EGFP cassette in the donor plasmid pBS-KS-attB1-2-PT-SA-SD-0-EGFP-FlAsHStrepII-TEV-3xFlag (Drosophila Genome Resource Center, Indiana University, Indiana, USA). For exchange, the donor plasmid was injected at a concentration of 50 ng/-l into eggs collected from a cross of *y^1^* M{vas-int.Dm}ZH-2A w^*^; M{3xP3-RFP.attP}ZH-86Fb virgin females with *betaTub97EF*^*MiMIC[MI06334]*^ males. Exchange events and associated cassette orientation were identified and characterized as described (Venken et al., 2011).

### Antibodies

For immunoblotting (IB) and immunofluorescence (IF) we used the following antibodies at the indicated dilutions: affinity purified rabbit anti-beta-Tubulin 97EF, 1:3000 (IF) and 1:8000 (IB); rabbit anti-beta-Tubulin 60D, 1:10000 (IF) and 1:5000 (IB); mouse monoclonal antibody DM1A against alpha-tubulin (Sigma, F2168) 1:10000 (IF) and 1:50000 (IB); mouse monoclonal anti-beta-tubulin TUB2.1 (Sigma, T4026) 1:500 (IB); mouse monoclonal antibody E-10 against beta-tubulin (Santa Cruz, sc-365791) 1:100 (IF) and 1:500 (IB); goat polyclonal antibody dN-17 against beta-tubulin (Santa Cruz, sc-20852), 1:500 (IB); mouse monoclonal antibody E7 against beta-tubulin 1:10 (IF) and 1:100 (IB); mouse monoclonal antibody Cq4 against Crumbs, 1:100 (IF); goat polyclonal antibody against GST (GE Healthcare Life Sciences, 27-4577-01), 1:1000 (IB); mouse monoclonal antibody anti-PSTAIR (Sigma Aldrich, P7962) 1:1000 (IB); mouse monoclonal antibody RL2 (Abcam, ab2739) 1:2000 (IB). Hybridoma supernatant containing E7 or Cq4 was obtained from Developmental Studies Hybridoma Bank (DSHB) created by the NICHD of the NIH and maintained at The University of Iowa, Department of Biology, Iowa City, IA 52242). Secondary antibodies for immunofluorescence were used at 1:500: Alexa 488-conjugated goat anti-rabbit IgG (Invitrogen, A-11008) and anti-mouse IgG (Invitrogen, A11029), Alexa 568-conjugated goat anti-rabbit IgG) (Invitrogen, A-11011) and anti-mouse IgG (Invitrogen, A-11004), or Cy5-conjugated goat anti-mouse IgG (Jackson ImmunoResearch, 115-175-146). Secondary antibodies used for immunoblotting (Jackson ImmunoResearch) were used at 1:1000: peroxidase-coupled goat anti-mouse and anti-rabbit, as well as well as donkey anti-goat.

### qRT-PCR

Total RNA was extracted from S2R+ cells (about 7x10^6^ cells) or embryos (100-300) using TRIzol (Invitrogen), followed by DNase digestion (DNA-Free Kit, Ambion). cDNA synthesis was performed using Transcriptor High-Fidelity cDNA Synthesis Kit (Roche) with 500ng RNA per reaction. Quantitative real-time PCR was performed using SYBR Green with an Applied Biosystems 7900HT using the recommended two-step cycling protocol. The following primer pairs (for sequences see Table S1) were used for the qRT-PCR analyses. For controls: *Act5C*: Act5C-F/Act5C-R, *Tbp*: Tbp-F/Tbp-R, *alphaTub84B*: alphaTub84B-F/alphaTub84B-R. For samples: *alphaTub84B*: FM158/159, *alphaTub84D*: FM160/161, *gammaTub23C*: FM176/FM177, *betaTub56D*: FM164/FM165, *betaTub60D*: FM170/FM171, *betaTub97EF*: FM106/FM107, FM79/80, FM63/FM66, FM39/FM40, FM64/FM65. Results obtained with the *betaTub97EF* primer pairs were averaged for Fig. 1. The *betaTub97EF* primer pairs used for Fig. 3 were FM63/FM66, FM73/FM74 and FM79/80. For distinction of the two *betaTub97EF* transcript isoforms resulting from mutually exclusive splicing of exon 4, the primer pairs FM63/FM66 (exon 4B) and FM63/FM111 (exon 4C) were used. For calculation of the ACT values (Fig. 1B), the CT values obtained for a given tubulin paralog was subtracted from an average CT value corresponding to the mean CT of the three control genes (*Act5C, alphaTub84B, Tbp*). We point out that our conclusions concerning relative abundance and temperature induced changes of the transcript levels of the tubulin paralogs based on qRT-PCR were not affected when only *Act5C* and *Tbp* but not *alphaTub84B* were used as control genes.

### Immunoblotting

Total extracts from bacteria expressing GST fusion proteins with N- and C-terminal extensions corresponding to those of the *Drosophila* beta-tubulin paralogs were made after transformation of *E. coli* BL21(DE3)[pREP4] with the pET-21d constructs described above. Expression was induced in transformants by addition of 1 mM IPTG during 2 hours. Cells were collected by centrifugation and lysed with 3x Laemmli buffer. The amount of GST fusion protein in the extracts was estimated after SDS-PAGE and Coomassie Blue staining gel using the staining intensity obtained with a known amount of molecular marker proteins resolved in parallel (PageRuler, Unstained Protein Ladder, Thermo Fisher Scientific, catalog number 26614).

For analysis of the expression of *Drosophila* beta-tubulin paralogs during embryogenesis, *w*^*1*^ and *betaTub97EF*^*MiMIC*^ embryos were collected for 3 hours and aged as indicated (Fig. 2A). Total extracts were prepared as described below. Ten embryos were loaded per lane. Two biological replicas gave similar results.

To compare the levels of beta-Tubulin 97EF protein in different genotypes, embryos were collected at 25°C from *w*^*1*^, *betaTub97EF*^*MiMIC*^ and *betaTub97EF*^*EGFP*^ and aged to 15-16 hours AED embryos. Total extracts were prepared as described below. 20 embryos were loaded per lane.

For analysis of the expression of *Drosophila* beta-tubulin paralogs during the larval stages, eggs were first collected at 25°C during one hour. The apple agar collection plate with the eggs was divided into five parts. These were aged at 25°C to either the late embryonic stage (20 hours AED), to the middle of first larval instar stage (36 hours AED), to the middle of the second (60 hours AED), and to the beginning or the end of the third larval instar stage (84 and 108 hours AED, respectively). Embryos at the late embryonic stage as well as the first instar larvae were collected directly from the plate into 0.7% NaCl, 0.07% Triton X-100 in an Eppendorf tube. Embryos were further processed as described (Radermacher et al., 2014). To obtain the other samples, the pieces of collection plate with eggs were placed into a bottle with standard fly food and aged for the desired time at 25°C. The upper layer of fly food with the larvae was then transferred into to a glass beaker, except for the late third instar samples. After addition of some water and a lid, the beaker was shaken to release the larvae from the food. After transfer of the beaker content into a glass dish, about 40 larvae were picked individually into an Eppendorf tube for sample preparation. Around 20 late third instar larvae were directly picked as wandering stage larvae from the walls of the bottles. Extracts were prepared as described below. 10-g of total protein was loaded per lane. Three biological replicas were performed.

To prepare samples from adults, 10 males and females (0-1 day after eclosion), respectively, were collected in an Eppendorf tube followed by extract preparation as described below. 10-g of total protein was loaded per lane.

Total extracts of *Drosophila* cells, embryos, larvae and adults were prepared in 3x Laemmli buffer. For extracts from larvae and adults, several inhibitors were added to the 3x Laemmli buffer: 1:20 Protease Inhibitor Cocktail, 1:100 PEFA Block 200 mM, Benzamidin 4 mM, Sigma). In general, samples were homogenized in 50-l and combined with additional 50-l used for washing homogenization pestle and tubes. Thereafter samples were heated for 5 minutes at 95°C, centrifuged for 5 minutes at maximum speed at 4°C in an Eppendorf centrifuge, aliquoted and snap frozen in liquid nitrogen. Total protein concentration in the extracts was determined using the Pierce 660nm Protein Assay (Thermo Scientific).

### Immunofluorescence

For high magnification analysis of the hindgut, *w*^*1*^ embryos (13-15 hours AED) were analyzed with an Olympus FluoView1000 confocal laser scanning microscope and a 60x/1.35 objective. For higher view of subcellular structures, a 2x zoom was used.

Third instar larvae and adult animals were dissected in phosphate buffered saline (PBS) and organs were directly transferred into 4% paraformaldehyd (PFA) in PBS for fixation. Samples were fixed for 30 min at room temperature (RT), followed by three 20 min washes with PBs containing 0.1% Triton X-100 (PBTx). Blocking was performed for 1 hour at RT in PBTx with 10% fetal bovine serum (FBS). Primary antibodies were added in blocking solution and incubated overnight at 4°C. After three 20 min washes with PBTx, the secondary antibodies were added in PBTx containing 10% FBS for 2 hours at RT. After three 20 min washes with PBTx, the tissues were incubated for 4 min at RT with Hoechst 33285 (1 μg/ml in PBTx). After three washes with PBS, tissues were mounted on slides in mounting media. For staining of ovaries, these were dissected from 3 day-old females. The ovarioles were gently separated apart partially with a dissection needle. Fixing, staining and imaging were performed as described above.

### Electron microscopy

Eggs were collected from *w*^*1*^ and *betaTub97EF*^*MiMIC*^ and aged at 25°C to 15-16 hours AED. Eggs were dechorionated. Unfertilized eggs were discarded. Embryos were prepared for transmission electron microscopy (TEM) and focused-ion beam scanning electron microscopy (FIBSEM). For high-pressure freezing, embryos were transferred from an agar plate to the 150-m cavity of a 3 mm aluminum specimen carrier using a needle or a pair of tweezers. 1-hexadecene was added on top and wicked off with a filter paper tip. Dextran (20% in PBS) was added on top to keep the embryos stuck to the carrier after freezing. The specimen carrier assembly was completed with a flat 3 mm aluminum specimen carrier lid. Samples were frozen in a HPM 100 high-pressure freezing machine (Leica Microsystems, Vienna, Austria).

For freeze-substitution and embedding the samples were sequentially incubated in water-free acetone with 1% OsO4 for 8 hours at −90°C, gradually warmed to −60°C (30°C/hour), for 6 hours at −60°C, gradually warmed to −30°C (30°C/hour), for 3 hours at −30°C, gradually warmed to 0°C (30°C/hour), 1 hour at 0°C. Subsequently, samples were washed twice with water-free acetone at 4°C, block stained with 1% uranyl acetate in water-free acetone for 1 hour at 4°C (uranyl acetate was diluted from a 20% stock solution in water free methanol), rinsed twice with water-free acetone and finally imbedded in Epon/Araldite (Sigma-Aldrich), 66% Epon/Araldite at 4°C overnight, 100% Epon/Araldite for 1 hour, followed by polymerization at 60°C for at least 48 hours. Ultra-thin sections were prepared with a Leica EM FCS ultra microtome (Leica Microsystems). TEM images were acquired with a FEI Tecnai G2 Spirit TEM at an acceleration voltage of 120 kV (Thermo Fisher Scientific, Eindhoven, NL) equipped with a Gatan Orius 1000 digital Camera (Gatan Inc., Pleasanton, California, USA).

For FIBSEM tomography, a trimmed Epon/Araldite block containing the sample was mounted on a scanning electron microscopy (SEM) stub using conductive carbon and coated with 10 nm of carbon by electron-beam evaporation to render the sample conductive. Ion milling and image acquisition was performed in an Auriga 40 Crossbeam system (Zeiss, Oberkochen, Germany) using the FIBICS Nanopatterning Engine (Fibics Inc., Ottawa, Canada). For each sample, a trench was milled at a current of 20 nA and 30 kV, followed by fine milling at 600 pA during image acquisition with a feed of 10 nm / 5 nm per image, respectively. Prior to starting the fine milling and imaging, a platinum layer of a few hundred nm was locally applied on top of the surface of the area of interest using the single gas injection system of the FIB-SEM. SEM images were acquired at 1.85 kV (60 μm aperture and high-current mode) using an energy-selective back-scattered electron detector with a grid voltage of 1.4 kV. The pixel size was 5 nm. Imaging was tilt-corrected to obtain square pixels. The images where aligned in TrakEM2 using SIFT (Cardona et al., 2012). Finally the resulting volumes were segmented semi-manually with the carving module of Ilastik (Andres et al., 2012; Straehle et al., 2011) and visualized with Imaris (Bitplane AG, Zurich, Switzerland).

**Table S1:**
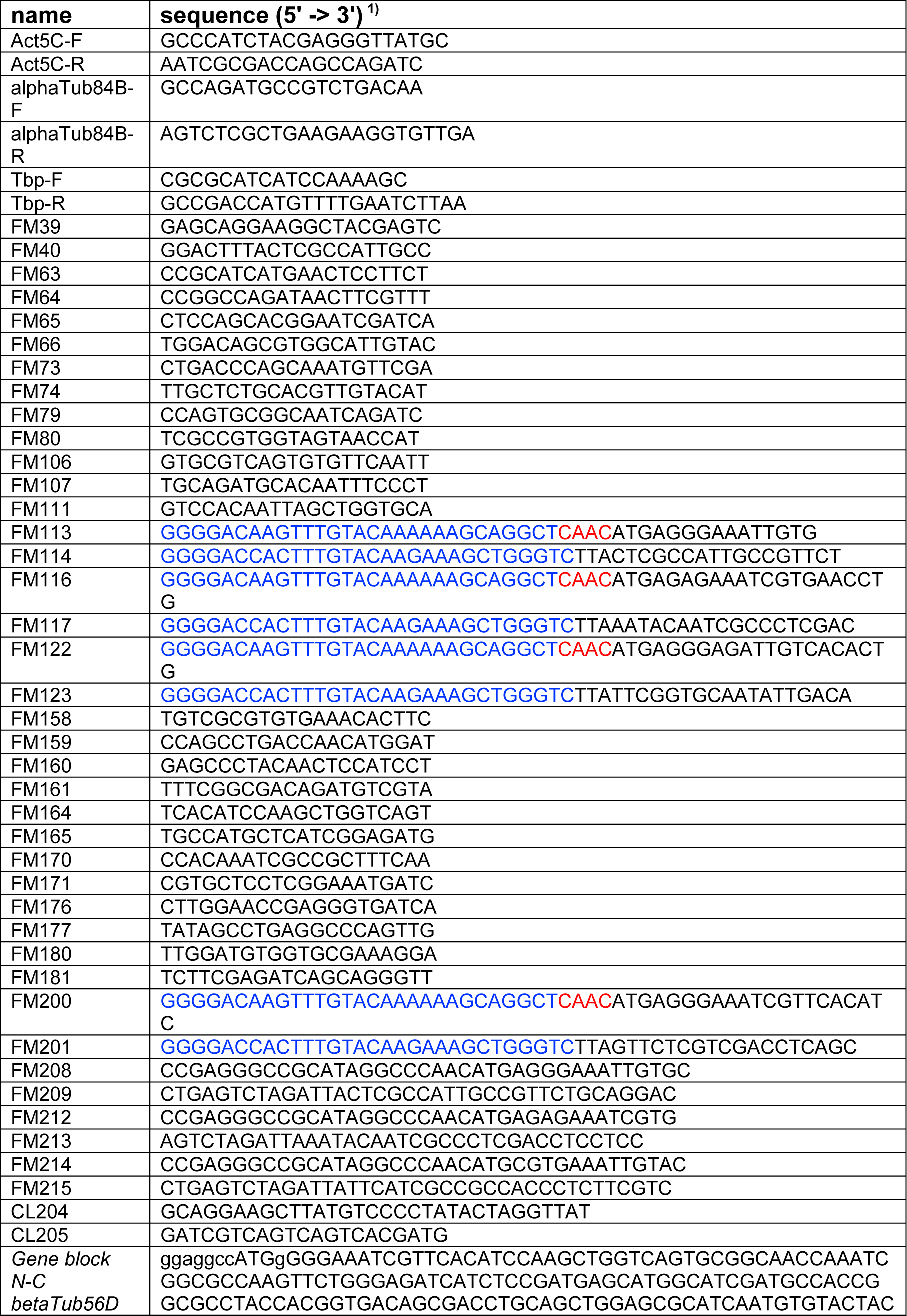
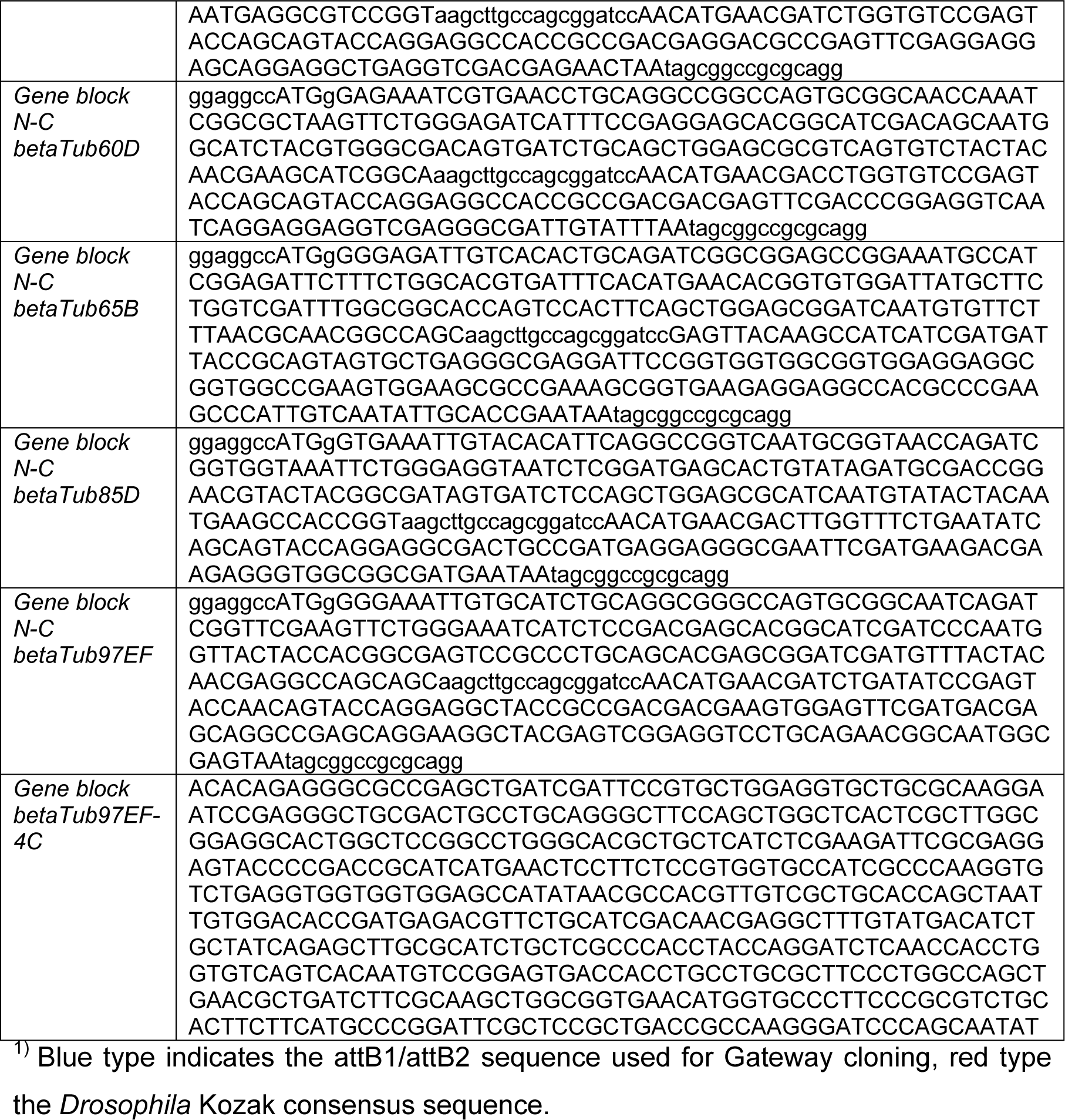
oligonucleotides and gene block sequences

## References

Althoff, F., Karess, R.E. and Lehner, C.F. (2012). Spindle checkpoint-independent inhibition of mitotic chromosome segregation by Drosophila Mps1. Mol Biol Cell 23, 2275–2291.

Bahi-Buisson, N., Poirier, K., Fourniol, F., Saillour, Y., Valence, S., Lebrun, N., Hully, M., Fallet Bianco, C., Boddaert, N., Elie, C., et al. (2014). The wide spectrum of tubulinopathies: what are the key features for the diagnosis? Brain 137, 1676–1700.

Bellen, H.J., Levis, R.W., Liao, G., He, Y., Carlson, J.W., Tsang, G., Evans-Holm, M., Hiesinger, P.R., Schulze, K.L., Rubin, G.M., et al. (2004). The BDGP gene disruption project: single transposon insertions associated with 40% of Drosophila genes. Genetics 167, 761–781.

Bischof, J., Bjorklund, M., Furger, E., Schertel, C., Taipale, J. and Basler, K. (2013). A versatile platform for creating a comprehensive UAS-ORFeome library in Drosophila. Development 140, 2434–2442.

Blose, S.H., Meltzer, D.I. and Feramisco, J.R. (1984). 10-nm filaments are induced to collapse in living cells microinjected with monoclonal and polyclonal antibodies against tubulin. The Journal of Cell Biology 98, 847–858.

Bossing, T., Barros, Claudia S., Fischer, B., Russell, S. and Shepherd, D. (2012). Disruption of Microtubule Integrity Initiates Mitosis during CNS Repair. Developmental Cell 23, 433–440.

Breitling, F. and Little, M. (1986). Carboxy-terminal regions on the surface of tubulin and microtubules epitope locations of YOL1/34, DM1A and DM1B. Journal of Molecular Biology 189, 367–370.

Buttgereit, D., Leiss, D., Michiels, F. and Renkawitz-Pohl, R. (1991). During Drosophila embryogenesis the beta 1 tubulin gene is specifically expressed in the nervous system and the apodemes. Mech Dev 33, 107–118.

Buttgereit, D., Paululat, A. and Renkawitz-Pohl, R. (1996). Muscle development and attachment to the epidermis is accompanied by expression of beta 3 and beta 1 tubulin isotypes, respectively. The international Journal of developmental Biology 40, 189–196.

Cherbas, L., Willingham, A., Zhang, D., Yang, L., Zou, Y., Eads, B.D., Carlson, J.W., Landolin, J.M., Kapranov, P., Dumais, J., et al. (2011). The transcriptional diversity of 25 Drosophila cell lines. Genome Research 21, 301–314.

Chiappori, F., Pucciarelli, S., Merelli, I., Ballarini, P., Miceli, C. and Milanesi, L. (2012). Structural thermal adaptation of beta-tubulins from the Antarctic psychrophilic protozoan Euplotes focardii. Proteins 80, 1154–1166.

Chu, D.T.W. and Klymkowsky, M.W. (1989). The appearance of acetylated α-tubulin during early development and cellular differentiation in Xenopus. Developmental Biology 136, 104–117.

Cleveland, D.W. (1988). Autoregulated instability of tubulin mRNAS: a novel eukaryotic regulatory mechanism. Trends in Biochemical Sciences 13, 339–343.

Cook, R.K., Christensen, S.J., Deal, J.A., Coburn, R.A., Deal, M.E., Gresens, J.M., Kaufman, T.C. and Cook, K.R. (2012). The generation of chromosomal deletions to provide extensive coverage and subdivision of the Drosophila melanogaster genome. Genome Biol 13, R21.

Correia, J.J. and Williams, J. R.C. (1983). Mechanisms of Assembly and Disassembly of Microtubules. Annual Review of Biophysics and Bioengineering 12, 211–235.

Coutelis, J.B., González-Morales, N., Géminard, C. and Noselli, S. (2014). Diversity and convergence in the mechanisms establishing L/R asymmetry in metazoa. EMBO reports 15, 926–937.

Delphin, C., Bouvier, D., Seggio, M., Couriol, E., Saoudi, Y., Denarier, E., Bosc, C., Valiron, O., Bisbal, M., Arnal, I., et al. (2012). MAP6-F is a temperature sensor that directly binds to and protects microtubules from cold-induced depolymerization. J Biol Chem 287, 35127–35138.

Denlinger, D.L. and Lee, R.E. (2010). Low temperature biology of insects. Cambridge: Cambridge University Press.

Detrich, H.W., 3rd, Parker, S.K., Williams, R.C., Jr., Nogales, E. and Downing, K.H. (2000). Cold adaptation of microtubule assembly and dynamics. Structural interpretation of primary sequence changes present in the alpha- and beta-tubulins of Antarctic fishes. J Biol Chem 275, 37038–37047.

Findeisen, P., Mühlhausen, S., Dempewolf, S., Hertzog, J., Zietlow, A., Carlomagno, T. and Kollmar, M. (2014). Six Subgroups and Extensive Recent Duplications Characterize the Evolution of the Eukaryotic Tubulin Protein Family. Genome Biology and Evolution 6, 2274–2288.

Fuller, M.T., Caulton, J.H., Hutchens, J.A., Kaufman, T.C. and Raff, E.C. (1988). Mutations that encode partially functional beta 2 tubulin subunits have different effects on structurally different microtubule arrays. The Journal of Cell Biology 107, 141–152.

Fygenson, D.K., Braun, E. and Libchaber, A. (1994). Phase diagram of microtubules. Physical Review E 50, 1579–1588.

Gadadhar, S., Bodakuntla, S., Natarajan, K. and Janke, C. (2017). The tubulin code at a glance. Journal of Cell Science 130, 1347–1353.

Gasch, A.P., Spellman, P.T., Kao, C.M., Carmel-Harel, O., Eisen, M.B., Storz, G., Botstein, D. and Brown, P.O. (2000). Genomic Expression Programs in the Response of Yeast Cells to Environmental Changes. Molecular Biology of the Cell 11, 4241–4257.

Graveley, B.R., Brooks, A.N., Carlson, J.W., Duff, M.O., Landolin, J.M., Yang, L., Artieri, C.G., van Baren, M.J., Boley, N., Booth, B.W., et al. (2011). The developmental transcriptome of Drosophila melanogaster. Nature 471, 473–479.

Gutzeit, H. (1986). The role of microtubules in the differentiation of ovarian follicles during vitellogenesis in Drosophila. Roux.Arch.Dev.Biol. 195, 173–181.

Hacker, U. and Perrimon, N. (1998). DRhoGEF2 encodes a member of the Dbl family of oncogenes and controls cell shape changes during gastrulation in Drosophila. Genes Dev 12, 274–284.

Hayashi, S., Ito, K., Sado, Y., Taniguchi, M., Akimoto, A., Takeuchi, H., Aigaki, T., Matsuzaki, F., Nakagoshi, H., Tanimura, T., et al. (2002). GETDB, a database compiling expression patterns and molecular locations of a collection of gal4 enhancer traps. Genesis 34, 58–61.

Hayward, S.A., Murray, P.A., Gracey, A.Y. and Cossins, A.R. (2007). Beyond the lipid hypothesis: mechanisms underlying phenotypic plasticity in inducible cold tolerance. Adv Exp Med Biol 594, 132–142.

Hazelett, D.J., Bourouis, M., Walldorf, U. and Treisman, J.E. (1998). decapentaplegic and wingless are regulated by eyes absent and eyegone and interact to direct the pattern of retinal differentiation in the eye disc. Development 125, 3741–3751.

Hoogenraad, C.C., Akhmanova, A., Grosveld, F., De Zeeuw, C.I. and Galjart, N. (2000). Functional analysis of CLIP-115 and its binding to microtubules. Journal of Cell Science 113, 2285–2297.

Hoyle, H.D. and Raff, E.C. (1990). Two Drosophila beta tubulin isoforms are not functionally equivalent. J. Cell Biol. 111, 1009–1026.

Hughes, J.R., Meireles, A.M., Fisher, K.H., Garcia, A., Antrobus, P.R., Wainman, A., Zitzmann, N., Deane, C., Ohkura, H. and Wakefield, J.G. (2008). A Microtubule Interactome: Complexes with Roles in Cell Cycle and Mitosis. PLOS Biology 6, e98.

Iwaki, D.D. and Lengyel, J.A. (2002). A Delta–Notch signaling border regulated by Engrailed/Invected repression specifies boundary cells in the Drosophila hindgut. Mechanisms of Development 114, 71–84.

Janke, C. (2014). The tubulin code: Molecular components, readout mechanisms, and functions. The Journal of Cell Biology 206, 461–472.

Jankovics, F. and Brunner, D. (2006). Transiently Reorganized Microtubules Are Essential for Zippering during Dorsal Closure in Drosophila melanogaster. Developmental Cell 11, 375–385.

Jaqaman, K., Loerke, D., Mettlen, M., Kuwata, H., Grinstein, S., Schmid, S.L. and Danuser, G. (2008). Robust single-particle tracking in live-cell time-lapse sequences. Nat Meth 5, 695–702.

Kemphues, K.J., Kaufman, T.C., Raff, R.A. and Raff, E.C. (1982). The testis-specific β-tubulin subunit in drosophila melanogaster has multiple functions in spermatogenesis. Cell 31, 655–670.

Kimble, M., Dettman, R.W. and Raff, E.C. (1990). The beta-3-tubulin gene of Drosophila-melanogaster is essential for viability and fertility. Genetics 126, 991–1005.

Kimble, M., Incardona, J. and Raff, E. (1989). A variant beta-tubulin isoform of Drosophila melanogaster (beta 3) is expressed primarily in tissues of mesodermal origin in embryos and pupae, and is utilized in populations of transient microtubules. Developmental Biology 131, 415–429.

Kumichel, A. and Knust, E. (2014). Apical Localisation of Crumbs in the Boundary Cells of the Drosophila Hindgut Is Independent of Its Canonical Interaction Partner Stardust. PLoS ONE 9, e94038.

Lee, T. and Luo, L. (1999). Mosaic Analysis with a Repressible Cell Marker for Studies of Gene Function in Neuronal Morphogenesis. Neuron 22, 451–461.

Leiss, D., Hinz, U., Gasch, A., Mertz, R. and Renkawitz-Pohl, R. (1988). Beta 3 tubulin expression characterizes the differentiating mesodermal germ layer during Drosophila embryogenesis. Development 104, 525–531.

Leśniewska, K., Warbrick, E. and Ohkura, H. (2014). Peptide aptamers define distinct EB1- and EB3-binding motifs and interfere with microtubule dynamics. Molecular Biology of the Cell 25, 1025–1036.

Lu, W., Winding, M., Lakonishok, M., Wildonger, J. and Gelfand, V.I. (2016). Microtubule– microtubule sliding by kinesin-1 is essential for normal cytoplasmic streaming in Drosophila oocytes. Proceedings of the National Academy of Sciences 113, E4995–E5004.

Ludueña, R. and Banerjee, A. (2008). The isotypes of tubulin. In The Role of Microtubules in Cell Biology, Neurobiology, and Oncology (ed. T. Fojo), pp. 123–175. New York, NY: Humana Press.

Metaxakis, A., Oehler, S., Klinakis, A. and Savakis, C. (2005). Minos as a genetic and genomic tool in Drosophila melanogaster. Genetics 171, 571–581.

Mimori-Kiyosue, Y., Shiina, N. and Tsukita, S. (2000). The dynamic behavior of the APC-binding protein EB1 on the distal ends of microtubules. Current Biology 10, 865–868.

Mitchison, T. and Kirschner, M. (1984). Dynamic instability of microtubule growth. Nature 312, 237–242.

Modig, C., Wallin, M. and Olsson, P.E. (2000). Expression of cold-adapted beta-tubulins confer cold-tolerance to human cellular microtubules. Biochem Biophys Res Commun 269, 787–791.

Molodtsov, M.I., Mieck, C., Dobbelaere, J., Dammermann, A., Westermann, S. and Vaziri, A. (2016). A Force-Induced Directional Switch of a Molecular Motor Enables Parallel Microtubule Bundle Formation. Cell 167, 539–552.e514.

Nogales, E. and Zhang, R. (2016). Visualizing Microtubule Structural Transitions and Interactions with Associated Proteins. Current opinion in structural biology 37, 90–96.

Pamula, M.C., Ti, S.-C. and Kapoor, T.M. (2016). The structured core of human - tubulin confers isotype-specific polymerization properties. The Journal of Cell Biology 213, 425–433.

Popodi, E.M., Hoyle, H.D., Turner, F.R. and Raff, E.C. (2008). Cooperativity beween the β-tubulin carboxy tail and the body of the molecule is required for microtubule function. Cell Motility and the Cytoskeleton 65, 955–963.

Radermacher, P.T., Myachina, F., Bosshardt, F., Pandey, R., Mariappa, D., Muller, H.A. and Lehner, C.F. (2014). O-GlcNAc reports ambient temperature and confers heat resistance on ectotherm development. Proc Natl Acad Sci U S A 111, 5592–5597.

Ravelli, R.B.G., Gigant, B., Curmi, P.A., Jourdain, I., Lachkar, S., Sobel, A. and Knossow, M. (2004). Insight into tubulin regulation from a complex with colchicine and a stathmin-like domain. Nature 428, 198–202.

Roth, S. and Lynch, J.A. (2009). Symmetry Breaking During Drosophila Oogenesis. Cold Spring Harbor perspectives in biology 1.

Rudolf, A., Buttgereit, D., Rexer, K.-H. and Renkawitz-Pohl, R. (2012). The syncytial visceral and somatic musculature develops independently of β3-Tubulin during Drosophila embryogenesis, while maternally supplied β1-Tubulin is stable until the early steps of myoblast fusion. European Journal of Cell Biology 91, 192–203.

Sampson, C.J. and Williams, M.J. (2012). Protocol for Ex Vivo Incubation of Drosophila Primary Post-embryonic Haemocytes for Real-Time Analyses. In Rho GTPases: Methods and Protocols (ed.F. Rivero), pp. 359–367. New York, NY: Springer New York.

Schulman, V.K., Folker, E.S. and Baylies, M.K. (2013). A method for reversible drug delivery to internal tissues of Drosophila embryos. Fly 7, 193–203.

Simcox, A., Mitra, S., Truesdell, S., Paul, L., Chen, T., Butchar, J.P. and Justiniano, S. (2008). Efficient genetic method for establishing Drosophila cell lines unlocks the potential to create lines of specific genotypes. PLoS Genet 4, e1000142.

Sirajuddin, M., Rice, L.M. and Vale, R.D. (2014). Regulation of microtubule motors by tubulin isotypes and post-translational modifications. Nat Cell Biol 16, 335–344.

Soplop, N.H., Cheng, Y.-S. and Kramer, S.G. (2012). Roundabout is required in the visceral mesoderm for proper microvillus length in the hindgut epithelium. Developmental Dynamics 241, 759–769.

Tartaglia, L.J. and Shain, D.H. (2008). Cold-adapted tubulins in the glacier ice worm, Mesenchytraeus solifugus. Gene 423, 135–141.

Taymaz-Nikerel, H., Cankorur-Cetinkaya, A. and Kirdar, B. (2016). Genome-Wide Transcriptional Response of Saccharomyces cerevisiae to Stress-Induced Perturbations. Frontiers in Bioengineering and Biotechnology 4, 17.

Ti, S.-C., Pamula, M.C., Howes, S.C., Duellberg, C., Cade, N.I., Kleiner, R.E., Forth, S., Surrey, T., Nogales, E. and Kapoor, T.M. (2016). Mutations in Human Tubulin Proximal to the Kinesin-Binding Site Alter Dynamic Instability at Microtubule Plus- and Minus-Ends. Developmental Cell 37, 72–84.

Valenstein, M.L. and Roll-Mecak, A. (2016). Graded Control of Microtubule Severing by Tubulin Glutamylation. Cell 164, 911–921.

Vemu, A., Atherton, J., Spector, J.O., Szyk, A., Moores, C.A. and Roll-Mecak, A. (2016). Structure and Dynamics of Single-isoform Recombinant Neuronal Human Tubulin. Journal of Biological Chemistry 291, 12907–12915.

Venken, K.J., Schulze, K.L., Haelterman, N.A., Pan, H., He, Y., Evans-Holm, M., Carlson, J.W., Levis, R.W., Spradling, A.C., Hoskins, R.A., et al. (2011). MiMIC: a highly versatile transposon insertion resource for engineering Drosophila melanogaster genes. Nat Methods 8, 737–743.

Wang, W., Zhang, H., Wang, X., Patterson, J., Winter, P., Graham, K., Ghosh, S., Lee, J.C., Katsetos, C.D., Mackey, J.R., et al. (2017). Novel mutations involving-I-,-IIA-, or-IVB-tubulin isotypes with functional resemblance to-III-tubulin in breast cancer. Protoplasma 254, 1163–1173.

Wheatley, S., Kulkarni, S. and Karess, R. (1995). Drosophila nonmuscle myosin-ii is required for rapid cytoplasmic transport during oogenesis and for axial nuclear migration in early embryos. Development 121, 1937–1946.

Widlund, P.O., Podolski, M., Reber, S., Alper, J., Storch, M., Hyman, A.A., Howard, J. and Drechsel, D.N. (2012). One-step purification of assembly-competent tubulin from diverse eukaryotic sources. Molecular Biology of the Cell 23, 4393–4401.

Wood, W. and Martin, P. (2017). Macrophage Functions in Tissue Patterning and Disease: New Insights from the Fly. Developmental Cell 40, 221–233.

Yu, I., Garnham, C.P. and Roll-Mecak, A. (2015). Writing and Reading the Tubulin Code. Journal of Biological Chemistry 290, 17163–17172.

Yu, N., Signorile, L., Basu, S., Ottema, S., Lebbink, J.H.G., Leslie, K., Smal, I., Dekkers, D., Demmers, J. and Galjart, N. (2016). Isolation of Functional Tubulin Dimers and of Tubulin-Associated Proteins from Mammalian Cells. Current Biology 26, 1728–1736.

Zhang, R., Alushin, G.M., Brown, A. and Nogales, E. (2015). Mechanistic Origin of Microtubule Dynamic Instability and Its Modulation by EB Proteins. Cell 162, 849–859.

## Supplementary References

Andres, B., Koethe, U., Kroeger, T., Helmstaedter, M., Briggman, K.L., Denk, W. and Hamprecht, F.A. (2012). 3D segmentation of SBFSEM images of neuropil by a graphical model over supervoxel boundaries. Med Image Anal 16, 796–796.

Bischof, J., Bjorklund, M., Furger, E., Schertel, C., Taipale, J. and Basler, K. (2013). A versatile platform for creating a comprehensive UAS-ORFeome library in Drosophila. Development 140, 2434–2434.

Bischof, J., Maeda, R.K., Hediger, M., Karch, F. and Basler, K. (2007). An optimized transgenesis system for Drosophila using germ-line-specific phiC31 integrases. Proc Natl Acad Sci U S A 104, 3312–3312.

Cardona, A., Saalfeld, S., Schindelin, J., Arganda-Carreras, I., Preibisch, S., Longair, M., Tomancak, P., Hartenstein, V. and Douglas, R.J. (2012). TrakEM2 software for neural circuit reconstruction. PLoS One 7, e38011.

Radermacher, P.T., Myachina, F., Bosshardt, F., Pandey, R., Mariappa, D., Muller, H.A. and Lehner, C.F. (2014). O-GlcNAc reports ambient temperature and confers heat resistance on ectotherm development. Proc Natl Acad Sci U S A 111, 5592–5592.

Rubin, G.M., Hong, L., Brokstein, P., Evans-Holm, M., Frise, E., Stapleton, M. and Harvey, D.A. (2000). A Drosophila complementary DNA resource. pp. 2222–2224.

Smith, D.B. and Johnson, K.S. (1988). Single-step purification of polypeptides expressed in Escherichia coli as fusions with glutathione S-transferase. Gene 67, 31–31.

Stapleton, M., Carlson, J., Brokstein, P., Yu, C., Champe, M., George, R., Guarin, H., Kronmiller, B., Pacleb, J., Park, S., et al. (2002). A Drosophila full-length cDNA resource. Genome Biol 3, RESEARCH0080.0081-0080.0088.

Straehle, C.N., Kothe, U., Knott, G. and Hamprecht, F.A. (2011). Carving: scalable interactive segmentation of neural volume electron microscopy images. Med Image Comput Comput Assist Interv 14, 653–653.

Venken, K.J., Schulze, K.L., Haelterman, N.A., Pan, H., He, Y., Evans-Holm, M., Carlson, J.W., Levis, R.W., Spradling, A.C., Hoskins, R.A., et al. (2011). MiMIC: a highly versatile transposon insertion resource for engineering Drosophila melanogaster genes. Nat Methods 8, 737–737.

